# Systematic comparison of sequencing-based spatial transcriptomic methods

**DOI:** 10.1101/2023.12.03.569744

**Authors:** Yue You, Yuting Fu, Lanxiang Li, Zhongming Zhang, Shikai Jia, Shihong Lu, Wenle Ren, Yifang Liu, Yang Xu, Xiaojing Liu, Fuqing Jiang, Guangdun Peng, Abhishek Sampath Kumar, Matthew E. Ritchie, Xiaodong Liu, Luyi Tian

**Author notes:** These authors contributed equally to this work (Yue You, Yuting Fu).

## Abstract

Recent advancements of sequencing-based spatial transcriptomics (sST) have catalyzed significant advancements by facilitating transcriptome-scale spatial gene expression measurement. Despite this progress, efforts to comprehensively benchmark different platforms are currently lacking. The extant variability across technologies and datasets poses challenges in formulating standardized evaluation metrics. In this study, we established a collection of reference tissues and regions characterized by well-defined histological architectures, and used them to generate data to compare six sST methods. We highlighted molecular diffusion as a variable parameter across different methods and tissues, significantly impacting the effective resolutions. Furthermore, we observed that spatial transcriptomic data demonstrate unique attributes beyond merely adding a spatial axis to single-cell data, including an enhanced ability to capture patterned rare cell states along with specific markers, albeit being influenced by multiple factors including sequencing depth and resolution. Our study assists biologists in sST platform selection, and helps foster a consensus on evaluation standards and establish a framework for future benchmarking efforts that can be used as a gold standard for the development and benchmarking of computational tools for spatial transcriptomic analysis.

## 1 Main

The advent of high-throughput sequencing technologies has revolutionized transcriptomics, providing unparalleled insights into the complexities of gene expression. Single-cell RNA sequencing (scRNA-seq) has been instrumental in dissecting cellular heterogeneity but falls short in capturing the spatial context essential for understanding tissue architecture, cellular interactions, and functional state [1, 2]. To address this limitation, sequencing-based spatial transcriptomics (sST) has emerged as a pivotal approach, enabling comprehensive transcriptomic profiling while preserving spatial information within tissues [3, 4].

Despite the rapid advancements in sST technologies, the field is still in its very early stages. The imaging-based spatial transcriptomics has a longer history and a collaborative benchmarking effort has been initiated with the SpaceTX consortium [5]. However, a systematic benchmarking study has not been done for sST. Prior studies have established frameworks for comparing single-cell transcriptomic and epigenomic methods, underscoring the necessity for standardized evaluation criteria and reference tissues for technology validation [6–9], since simulated single-cell and spatial data may not be reliable [10]. While sST technologies share common features, such as the use of spatial DNA barcodes analogous to cell barcodes in scRNA-seq, the methods diverge significantly in aspects like spatial resolution and the preparation of spatially barcoded oligo arrays [11]. This variability introduces challenges in method selection and complicates the establishment of universal evaluation standards.

In the present study, we address this critical gap by conducting a systematic comparison of six sST methods. Using a set of reference tissues, including mouse embryonic eyes and hippocampal regions of the mouse brain, we generated cross-platform data for sequencing-based ST benchmarking, referred to as *cadasSTre*. This dataset enables us to evaluate the performance of each technology in terms of spatial resolution, capture efficiency, and molecular diffusion. We updated *scPipe* [12] to enable preprocessing and downsampling of sST data, to further minimize variability and facilitate the incorporation of future technologies. Our analyses reveal that data generated from different sST technologies exhibit varying capabilities in downstream applications, such as clustering, region annotation, and cell-cell communication. Notably, we also highlighted gene detection biases in sST data.

Our study serves multiple purposes: it (i) guides researchers in the selection of appropriate sST methods for their specific biological questions, (ii) establishes a framework for future benchmarking endeavors, and (iii) contributes to the standardization of evaluation criteria in this rapidly evolving field. Furthermore, our work aims to provide a foundation for the assessment of computational tools designed for spatial transcriptomic data analysis.

## 2 Results

### 2.1 Benchmarking reference tissues and experimental design

We systematically benchmarked spatial transcriptomics (sST) methods based on distinct spatial indexing strategies, encompassing microarray (10X Genomics Visium [13]), bead-based approaches (HDST [14], BMKMANU S1000, Slide-seq [15]), polony- or nanoball-based technologies (Stereo-seq [16], PIXEL-seq [17]), and microfluidics (DBiT-seq [18]). Details of each sST method are listed in Supplementary Table 1.

**Table 1.**
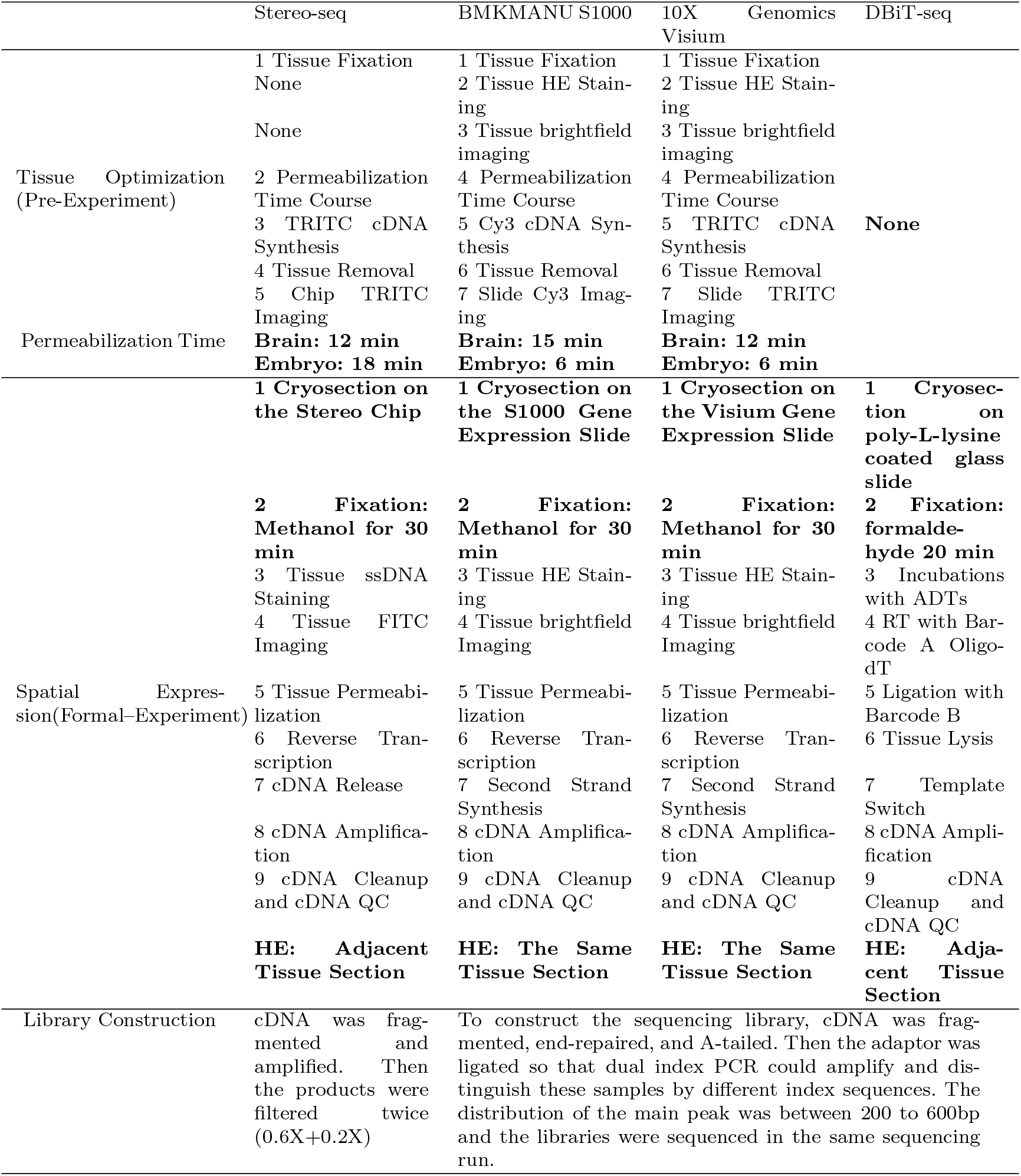
Protocols used in different ST methods.

We selected the adult mouse brain, E12.5 mouse embryo, and adult mouse olfactory bulb (OB) as reference tissues due to their relatively well-defined morphological characteristics. Adult mouse hippocampus, for instance, exhibits consistent thickness and comprises regions such as Cornu Ammonis (CA)1, CA2, CA3, and Dentate Gyrus (DG), each with distinct expression profiles. E12.5 mouse eyes in embryo exhibit a known structure with a lens surrounded by neuronal retina cells, while mouse olfactory bulbs (OB) feature clear layer separation with various neuron types. These tissues, with their known morphological patterns and heterogeneous expression profile, serve as ideal reference samples for our sST benchmark studies. The use of diverse tissue types allowed us to assess how tissue type influences method performance, and each sample included a technical replicate for variability assessment (Figure 1a). A summary of the datasets in *cadasSTre* is given in Supplementary Table 2. Detailed protocols for obtaining regions of interest have been established and are available in the Methods section, facilitating reproducibility by other researchers. In total, we systematically evaluated 6 sST methods across 22 experiments from 3 tissue types.

**Fig. 1.**
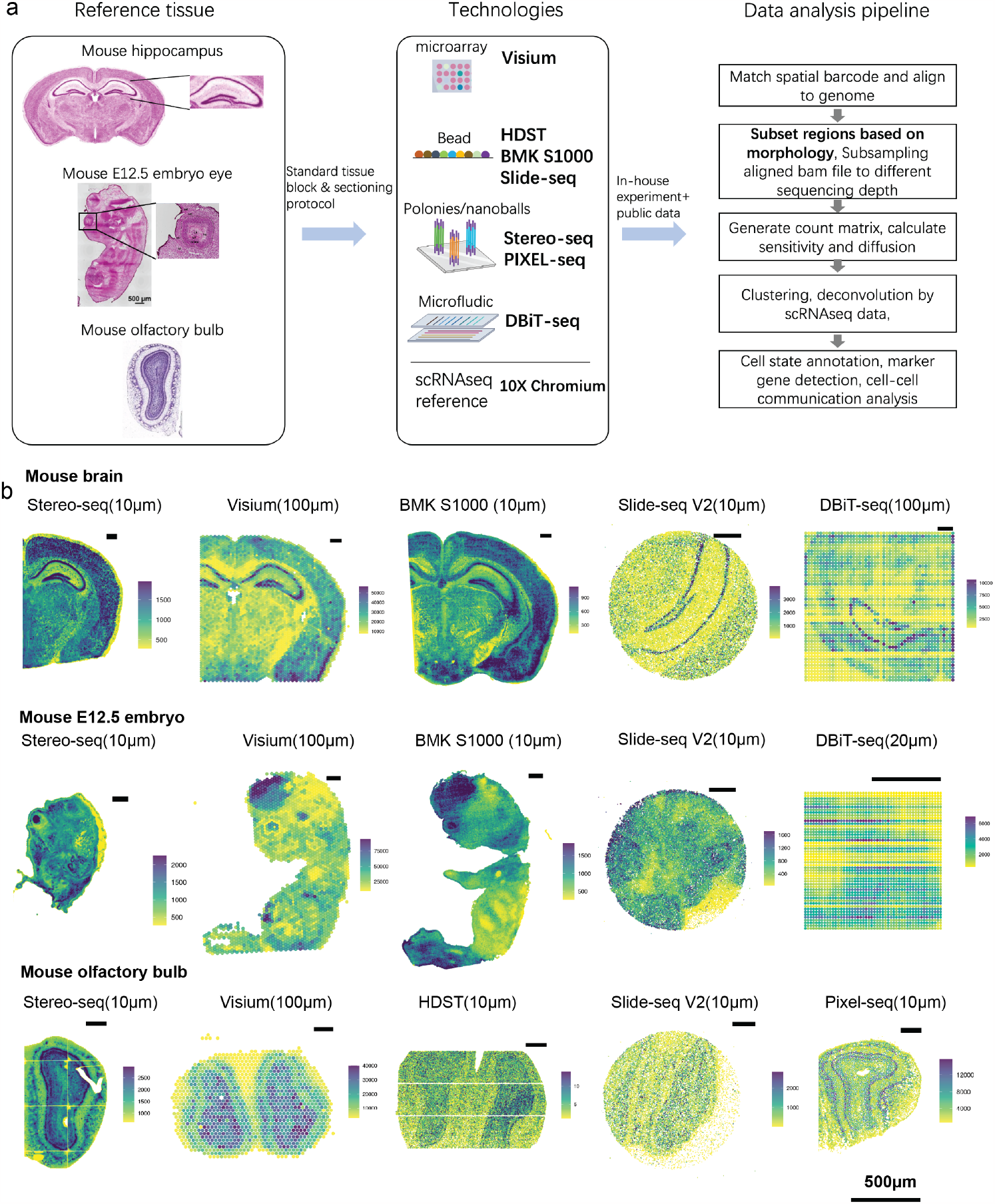
Overview of experimental design and data processing pipeline. **a)** The experimental design involved the use of reference tissues, namely, adult mouse hippocampus, E12.5 mouse eye, and adult mouse olfactory bulb. We performed sST on these reference tissues using diverse technologies categorized by their distinct spatial indexing strategies. These techniques encompassed microarray-based methods (e.g., 10X Genomics Visium), bead-based approaches (such as HDST, BMKMANU S1000 (abbreviation: BMK S1000), and Slide-seq), polonies or nanoballs techniques (Stereo-seq and PIXEL-Seq), and microfluidic-based methodologies like DBiT-seq. Additionally, the reference tissues were subjected to single-nuclei RNA-sequencing (snRNA-seq) using the 10X platform. The *cadasSTre* datasets underwent a series of processing steps. Initially, spatial barcodes, their corresponding locations, and expression profiles were generated. Subsequently, reads within regions with known morphology were selectively retained, and downsampling was performed to mitigate the impact of sequencing depth variations. Count matrices were then generated for sensitivity and diffusion calculations. This was followed by cell state annotation and a comprehensive analysis of cell-to-cell communication. **b)** The visualization of total counts across the spatial dimension for datasets generated using each platform for reference tissues is shown. The distances from center to center, used in creating the plot, are presented alongside the name of each sST method. The length of the black bar in the visualization corresponds to a distance of 500 microns.

As outlined in the summary pipeline (Figure 1a right-hand panel), we next built a standard benchmarking pipeline to enable homogeneous data processing for sST methods and comparison in a fair way. Initially, spatial barcodes and their corresponding locations, together with expression profiles per spatial location were generated. Figure 1b provides an overview of total counts per spot for each sST method across various tissue types. Clear tissue patterns were observed across the samples. The summary of total counts is presented with varying spot sizes and the distances between spot centers. These differences are clearly depicted in Supplementary Figure 1a. They exhibit clear differences, as depicted in Supplementary Figure 1a. In Figure 1b, we have labeled the distances between spot centers, as we believe this metric better represents the platform’s physical resolution, as opposed to using spot sizes. Stereo-seq and BMKMANU S1000 have distances between spot centers smaller than 10*µ*m and spots in them are binned into a 10*µ*m-sized spots for visualization.

We observed that Stereo-seq, Visium, and BMKMANU S1000 managed to capture nearly the entire right brain and the whole E12.5 embryo. In contrast, Slide-seq V2 could capture only a portion of the tissue due to its limited capture size (Supplementary Figure 1b,c). With DBiT-seq, the capture size varies depending on the width of the microfluidic channel, while also posing the risk of contamination across columns and rows in channels. We observed highly consistent tissue morphology among different methods in the H&E image shown in supplementary figure 2-4, which validates that our standard tissue handling and sectioning protocol could generate consistent results in different experiments.

**Fig. 2.**
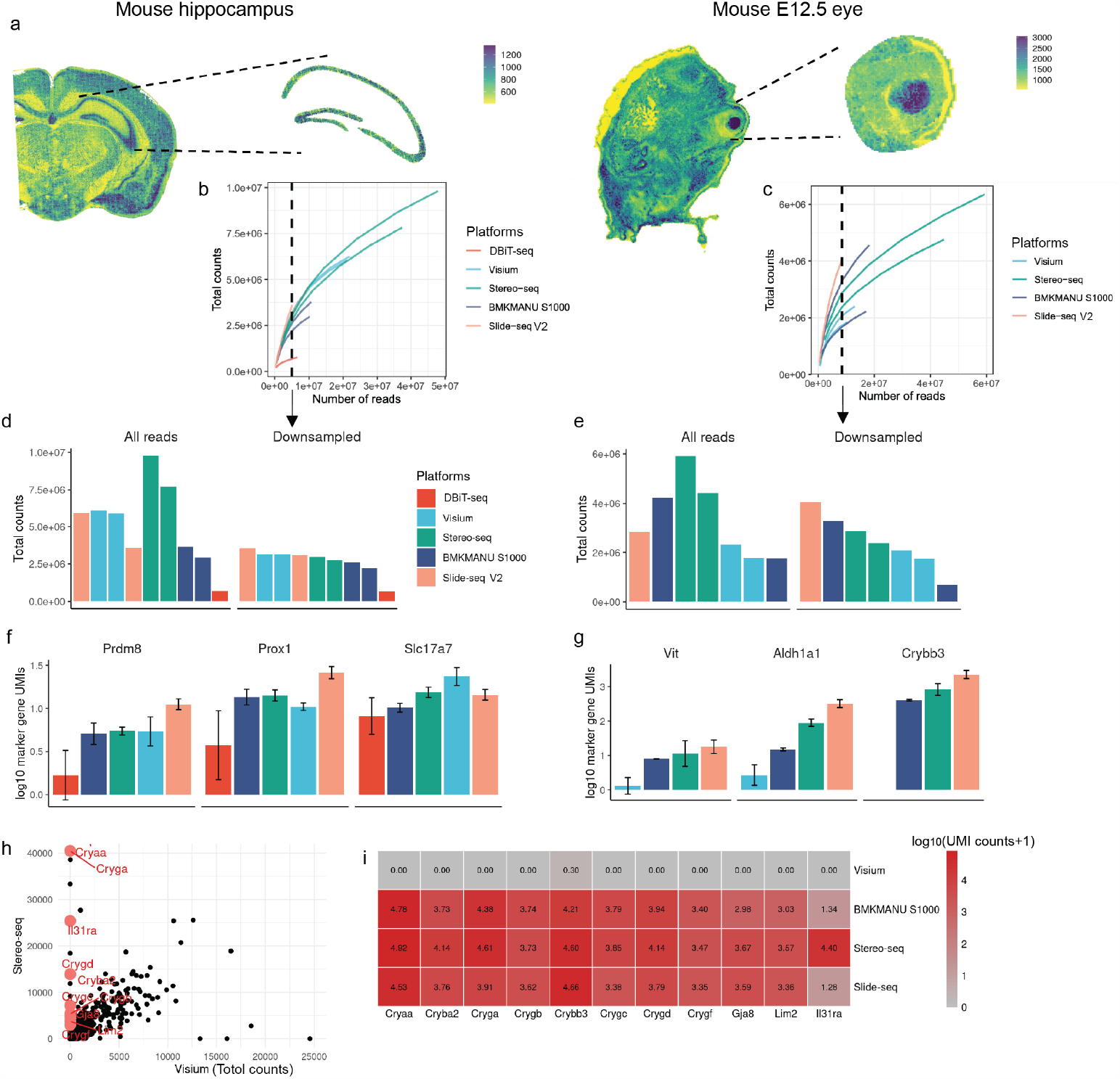
Comparison of the sensitivity of data generated by different platforms. **a)** Schematic plot illustrating the extraction of regions with known morphology from fully processed samples of the adult mouse hippocampus and E12.5 mouse eye. Total UMI counts are presented as a function of stepwise downsampled sequencing depths for each platform. The data originates from **b)** mouse hippocampus and **c)** E12.5 mouse eye regions. A vertical dashed black line marks the read count used for generating the subsequent downsampled data. **d)** Total unique molecular identifier (UMI) counts were computed for selected regions using all reads and downsampled data for the mouse hippocampus. **e)** Total UMI counts for selected regions using all reads and downsampled data for the E12.5 mouse eye. **f)** The summed UMI counts for marker genes across five individual 100*µ*m *×* 100*µ*m regions in the mouse hippocampus, along with mean and standard deviation. **g)** The summed UMI counts for marker genes across five individual 100*µ*m *×* 100*µ*m regions in the E12.5 mouse eye, along with mean and standard deviation. **h)** Total UMI counts of detected genes are compared between Visium (x-axis) and Stereoseq (y-axis). Each dot represents a gene, shown in black. Genes that display expression at the 90th percentile with Stereo-seq but are at the 10th percentile in Visium are highlighted in red and labeled with their gene symbols. **i)** A heatmap displays the log_10_-transformed expression of genes that are specifically not captured by Visium but are captured by Stereo-seq for E12.5 mouse eyes.

Subsequently, we selectively retained reads within regions with known morphology, including the hippocampus in the mouse brain, and eyes in the E12.5 embryo. We then performed downsampling to address sequencing depth and sequencing cost variations. The purpose of downsampling is to normalize different methods to the same total number of sequencing reads to achieve equivalence in sequencing cost. Count matrices with downsampled data and full data were both then generated for sensitivity and diffusion calculations, followed by cell state annotation, maker gene detection, and analysis of cell-to-cell communication.

### 2.2 Molecule-capture efficiency

We obtained hippocampus and eye tissues from the adult mouse brain and E12.5 mouse embryo, as illustrated in Figure 2a. This was accomplished by manually delineating boundaries based on tissue patterns indicated by the spatial distribution of total counts and morphological information provided by H&E images. By selecting the same region, we ensure that our comparisons of sST sample performance were not influenced by varying locations within the tissues, as the number of counts from different parts of the tissue may exhibit variations.

Molecule capture efficiency was assessed in two ways. In selected regions, we either 1) used all the reads from that region, or 2) downsampled the data so that different samples had the same number of sequenced reads, which we refer to as *“downsampled data*” in the subsequent results.

Based on the downsampling results (Figure 2b,c), none of the sequencing runs, that ranged from 300 million reads (Visium) to 4 billion reads (Stereo-seq), reached 5 saturation. This observation suggests that sST data requires considerably more reads for optimal performance, with the potential for increased sensitivity.

Next, we compared the sensitivity of each sST method by summing the total counts within the selected regions. Stereo-seq had many more sequencing reads for the same region compared to other platforms, resulting in higher total counts when all reads are used (Figure 2d and e, left panel). However, when the effect of sequencing depth is controlled, Slide-seq V2 data consistently demonstrated higher sensitivity than other platforms, in both the eye and hippocampus. This observation aligns with the saturation plot results (Figure 2b,c), where the total counts from Slide-seq V2 data exhibited a greater increase with increasing read number. In contrast, DBiT-seq data consistently showed the lowest sensitivity (Figure 2d and e, right panel). Additionally, the impact on the relationship between the number of counts and features per spot is more pronounced in Stereo-seq data when comparing downsampled results to the results obtained using all reads (Supplementary Figure 5).

To provide a more detailed assessment of the differences in sensitivity among selected sST methods, we proceeded to measure the RNA content of marker genes known to be expressed in specific regions using downsampled data. In CA3 of the hippocampus, we compared the sum of counts for *Prdm8, Prox1*, and *Slc17a7* within 100*µ*m *×* 100*µ*m regions (selected based on the largest physical resolution value among the sST methods applied). Our findings revealed that the expression patterns of these marker genes mirrored the total count results, with Slide-seq exhibiting the highest sensitivity and DBiT-seq displaying the lowest (Figure 2f). In the case of E12.5 mouse eyes, we compared the sum of counts for *Vit, Crybb3* (lens), and *Aldh1a1* (neuron retina) within 100*µ*m *×* 100*µ*m regions. Similarly, Slide-seq demonstrated the highest sensitivity, while Visium did not generate as many counts for marker genes in regions where their expression was expected (Figure 2g). Furthermore, through pairwise comparisons, we identified genes consistently expressed in the lens across all sST methods, except for data generated by Visium (Figure 2h, Supplementary Figure 6a), including *Crybb3* and *Cryaa* (Figure 2i). Importantly, this inconsistency did not appear to be attributed to the preprocessing pipeline and gene annotations (Supplementary Figure 6b), indicating a systematic gene-specific bias of Visium towards the lens. In an attempt to correlate this bias with various gene attributes, including gene biotypes, length, and GC content percentage, we discovered that these biased genes, which exhibit low expression in Visium, are predominantly protein-coding. Moreover, no significant bias was detected in terms of GC content or gene length (Supplementary Figure 7).

In our investigation of the mouse OB, after annotation, we assessed the sensitivity of selected sST methods, considering layers with varying densities of total counts. Notably, PIXEL-seq exhibited the highest sensitivity, while HDST demonstrated the lowest sensitivity at a 10*µ*m physical resolution (Supplementary Figure 8).

### 2.3 Molecule-lateral diffusion

In addition to molecule capture sensitivity per unit area, another crucial quality parameter is the spatial accuracy of mRNA detection. To assess such accuracy, we employed two analysis methods to measure molecule lateral diffusion: 1) Plotting the intensity profile of a specific gene across the selected region. 2) Quantifying the distance between the left width at half-maximum (LWHM) of intensity in the chosen region [19], focusing on histological structures where the expression of the selected gene should exhibit a significant difference—showing high expression in one part of the region and minimal to no expression in the rest. These analyses were conducted using count data generated from all reads.

In our evaluation of the OB, we selected *Slc17a7* as the marker gene due to its expected expression specifically in Mitral and Tufted (M/T) cells, which form distinct layers [20] and in glutamatergic neurons located at the base of the glomerular layer (GL) [21]. We confirmed *Slc17a7*’s expression at these locations via in situ hybridization (ISH) from the Allen Brain Atlas [22]. In this analysis, our focus was on *Slc17a7*’s expression in M/T cells. As illustrated by the expression plots of *Slc17a7* in each sST dataset (Figure 3a, left panel, Supplementary Figure 9), we specifically selected regions (N=6) where *Slc17a7* was expressed in the middle. Our observations, based on intensity plots and LWHM measurements, revealed significant lateral diffusion by Stereo-seq V1 of *Slc17a7* in the OB. Notably, Slide-seq V1.5 and PIXEL-seq exhibited relatively better control over this diffusion (Figure 3b-d left panel).

**Fig. 3.**
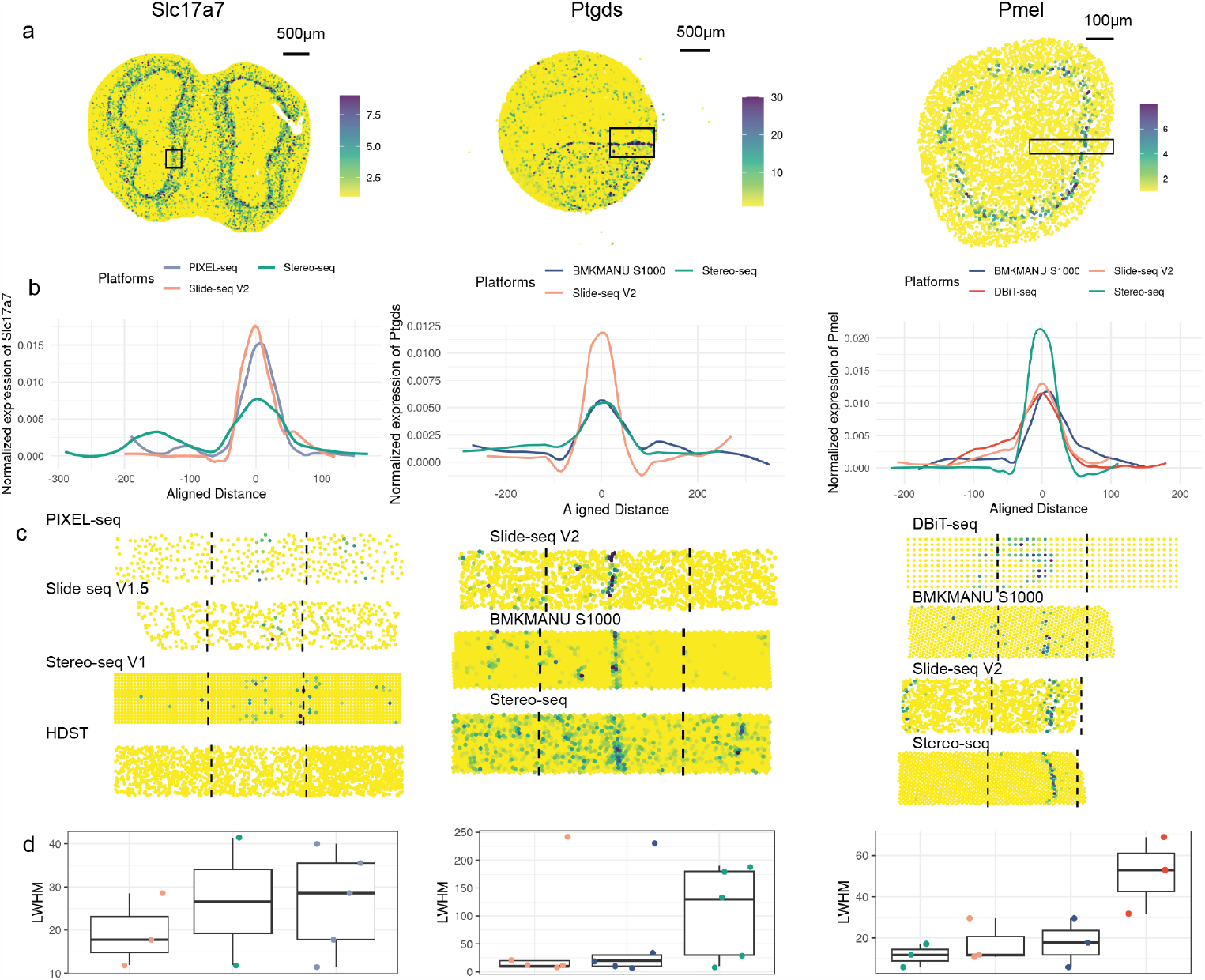
Comparison of diffusion of data generated by different platforms. **a)** Expression patterns of selected marker genes known to be highly expressed in specific regions. These markers include *Slc17a7* in the mouse olfactory bulb (left panel), *Ptgds* in the mouse brain (middle panel), and *Pmel* in the E12.5 eye (right panel). The plots are based on raw count values. Black boxes indicate the selected regions used for diffusion calculation. **b)** Expression levels of the aforementioned marker genes (from panel a) are aggregated for every 10µm along 50*µ*m in the olfactory bulb, 500*µ*m in the brain, and 300*µ*m in the eyes, as shown in a). UMI counts are averaged across modalities, normalized for each platform, and presented in a density plot with the area under the curve set to 1 (details in Methods). **c)** Expression level of the marker genes as mentioned above (from panel a) within selected modalities are provided, with black dashed lines delineating the boundaries used for diffusion calculations. **d)** The left half-width half maximum (LWHM) of the profile was then calculated for each gene (from panel a) in each modality and displayed in boxplots. Each dot represents the LWHM for a given modality. Modalities for which LWHM could not be calculated were removed.

In our analysis of the brain, we selected *Ptgds* as the marker gene, as it has been confirmed by ISH to be specifically expressed in a particular location within vascular cells [23] (Supplementary Figure 10a,b). By examining the expression plots of *Ptgds* and its intensity plots along with LWHM measurements, we noted severe lateral diffusion in the Stereo-seq dataset. In contrast, Slide-seq V2, followed by BMKMANU S1000, exhibited better control over such lateral diffusion issues (Figure 3a-d middle panel, Supplementary Figure 10c). We further validated these observations by conducting a diffusion analysis on downsampled Stereo-seq data, confirming that the challenge of lateral diffusion persisted despite a lower sequencing depth compared to other sST datasets. (Supplementary Figure 10d-f) This suggests that downsampling could not resolve the lateral diffusion issue for Stereo-seq data.

For our examination of eye tissue, we selected *Pmel* as the marker gene due to its specific expression in melanocytes, which encircle the lens and form a circular pattern [24]. Interestingly, in this context, Stereo-seq demonstrated the best control over lateral diffusion, followed by Slide-seq V2. (Figure 3a-d right panel, Supplementary Figure 11) This observation contrasts with our findings in the other two tissue types, indicating that tissue type exerts a considerable influence on the diffusion process. Diffusion is greatly impacted by permeabilization time. We have showed in our permeabilization optimization experiment (Supplementary Figure 2B,3B,4B) that different permeabilization time significant impact the diffusions.

### 2.4 Clustering and cell annotation across technologies

We next applied selected sST methods to gain insight into biological questions where higher capture sensitivity and well-controlled diffusion are important.

We selected E12.5 mice eyes, known for their distinctive structure featuring the lens, surrounded by the retina, and then melanocytes [25–27].

#### 2.4.1 Annotating regions by clustering results

With the basic knowledge of general cell states within the eye area, our next objective was to annotate the spots captured by selected sST platforms using various clustering methods. We aimed to determine whether we could consistently identify more detailed and coherent cell subsets across all samples.

Such resulting annotations of cell subsets not only served as a benchmark for evaluating the methods employed in this study but also provided valuable insights into the intricacies of cell states within the developing eye of E12.5 mice.

Before delving into our comparative analyses, Figure 4a showcases our findings about the cell subsets that we expected to observe within an E12.5 mouse eye. In this tissue, the anticipated morphological structure unfolds from the innermost space, housing the lens and lens vesicle, which are enveloped by neuronal retina cells forming distinct subsets in specific locations. The neuronal retina cells are encircled by melanocytes, with the rostral side hosting corneal mesenchyme, while the caudal side is composed of epithelial cells. These annotations provided us with a foundation for our subsequent evaluations and comparative assessments.

**Fig. 4.**
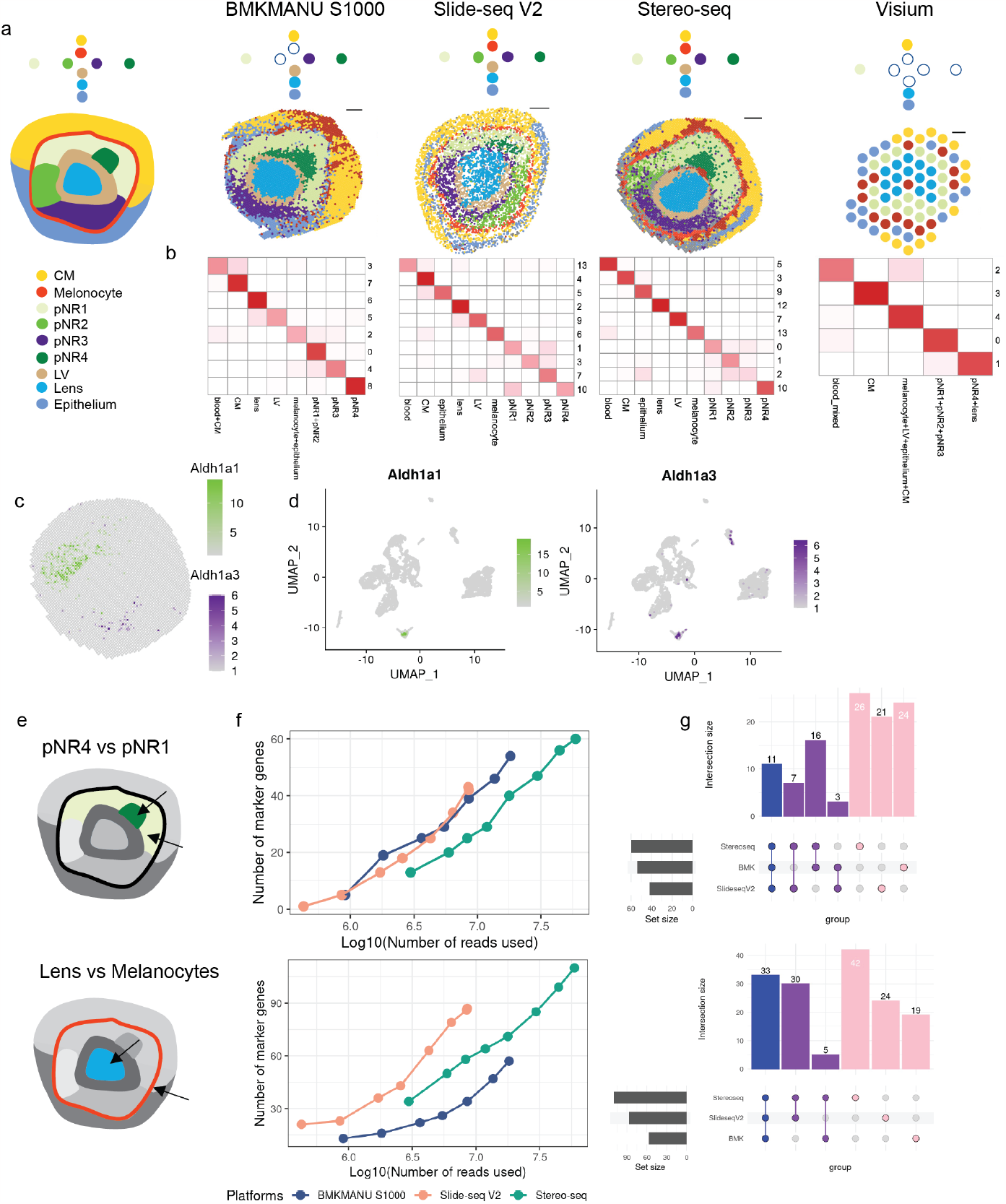
Comparison on downstream performance. **a)** Expression profiles generated by each platform were processed to obtain clustering results. Known cell types and states are colored in the left-most panel. Additionally, a schematic plot represents the expected cell states, arranged from outer space to inner space and from top to bottom. On the right-hand side, clustering results are presented, with spots color-coded by annotated cell states depicting the identifiable cell states. CM represents corneal mesenchyme; pNR represents presumptive neural retina; LV represents lens vesicle. **b)** Clustering was conducted on downsampled eye data from each platform, with an equal total read count across platforms in the eye area. The correspondence between annotations obtained from clustering based on all reads and clustering based on downsampled data is visualized in a heatmap. The number of spots in this correspondence is presented after log_10_ transformation without scaling. **c)** With Stereoseq data as an example, spatial expression profiles of *Aldh1a1* and *Aldh1a3*, which are expressed in pNR2 and pNR3 are shown at 10 *µ*m resolution. The number of spots with the expression of *Aldh1a1* above 0 is 1,329, and of *Aldh1a3* above 0 is 217. **d)** Expression profiles of *Aldh1a1* and *Aldh1a3* in snRNA-seq data are presented, The number of cells having an expression of *Aldh1a1* above 0 is 93, and of *Aldh1a3* above 0 is 161. **e)** An overview of cell states compared in the marker gene detection analysis, with pNR4 and pNR1 highlighted in the top panel, and lens and melanocytes highlighted in the bottom panel. **f)** Number of marker genes detected with different numbers of reads used for each sST method in the comparison between pNR4 and pNR1 (top panel). The same analysis is applied to the lens and melanocytes in the bottom panel. **g)** In the top panel, an Upset plot displays the intersection of marker genes obtained by different sST methods using all reads for the pNR4 and pNR1 comparison. Genes shared among all three platforms are denoted in blue, those shared between two platforms are in purple, and uniquely obtained genes are represented in pink. The bottom panel presents a similar analysis for lens and melanocytes.

**Fig. 5.**
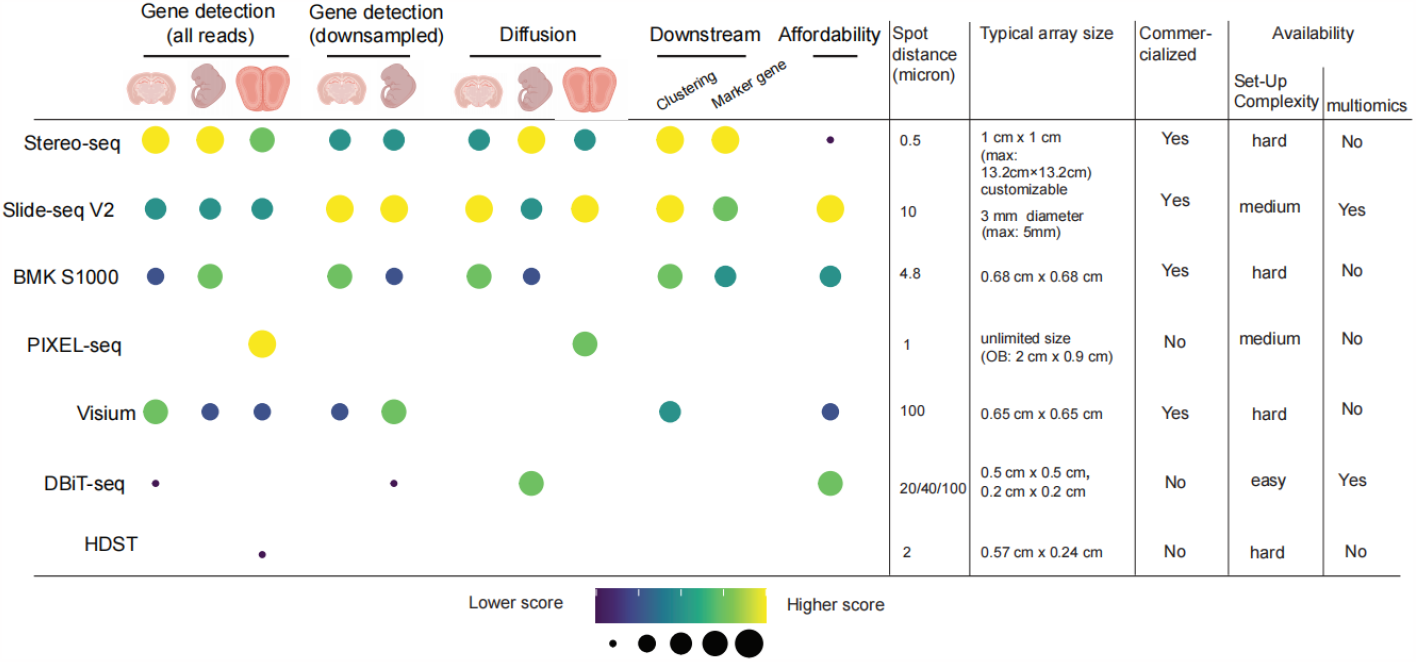
Summary of results and characteristics of sST methods. The sST methods have been ranked based on their performance in the specified categories, with the highest-performing methods positioned at the top. In the left panel, each ranking is represented by color and spot size. In the right panel, essential characteristics of the sST methods examined are outlined. Set-up complexity represent how difficult it is to build the method from scratch.

#### 2.4.2 Comparison between clustering results

In our comparative analysis of clustering results, we conducted evaluations from two perspectives:

Clustering Methods: We systematically employed three distinct clustering methods: *Seurat* [28], which exclusively considers transcriptomic profiles, and *DR*.*SC* [29] and *PRECAST* [30], which incorporate spatial information alongside gene expression data. Recent benchmark studies have reported that methods leveraging spatial location information demonstrate promising clustering results in specific datasets. However, they do not consistently surpass or exhibit greater robustness compared to methods that solely rely on gene expression data [31]. Our observations align with this conclusion, with *Seurat* consistently demonstrating robust and stable performance compared to the other 2 methods in detecting expected cell subsets as shown in Figure 4a, left panel, and Supplementary Figure 12.
sST Methods: In our comparisons between sST platforms, we focused primarily on the results generated by *Seurat*. We annotated spots for each sST method individually (Figure 4a and Supplementary Figure 13). Our analyses unveiled variations in the ability of different methods to consistently identify the expected cell subsets. Notably, Slide-seq V2 and Stereo-seq data delivered a nice separation of spots for comprehensive subset annotations, successfully capturing all anticipated subsets. Conversely, BMKMANU S1000 data faced challenges in cell state detection, particularly in identifying melanocytes. This difficulty may stem from the pronounced lateral diffusion observed in BMKMANU S1000 data (as depicted in Figure 3c,d right panel), making it difficult for clustering methods relying solely on expression profiles to retain this specific cell type. On the other hand, Visium data faced certain limitations in detecting the anticipated cell subsets. These challenges were primarily attributed to the relatively low physical resolution and hence a restricted number of spots available in the eye area (approximately 75 spots in total for each sample). Within each of these 100*µ*m *×* 100*µ*m spots, cells were mixed, making identifying intricate cell subsets more challenging (Figure 4a, right panel).

#### 2.4.3 Influence of downsampling on clustering results

We observed that sequencing depth influences the total counts of spatial transcriptomic data (Figure 2b-e). In light of this, we set out to investigate how sequencing depth impacts clustering results. Our exploration of clustering results on downsampled data involved two key aspects: 1) We assessed the correspondence between the downsampled data and the full data. 2) We calculated entropy measures for cluster purity (ECP) and accuracy (ECA) based on the clustering results obtained with the full data as a reference for downsampled data generated at various proportions as shown in Figure 4b. Remarkably, we discovered that the downsampled data was capable of detecting nearly all of the cell subsets identified by the full data (Figure 4b, Supplementary Figure 14). However, when evaluating ECP and ECA across different proportion values, we observed relatively high values, signifying a notable degree of inconsistency. This inconsistency could be attributed to the fact that while the majority of cell subsets effectively formed distinct clusters, a portion of cells grouped into different clusters, notably between cells from different subsets of neuronal retina cells. This effect was particularly pronounced in subsets between populations such as lens and lens vesicles; 4 neuron retina subsets (Figure 4b), which are more similar in expression profiles.

#### 2.4.4 Comparison between sST data and snRNA-seq data

We consistently observed well-patterned expression of *Pmel, Crybb3, Atoh7, Enfa5, Aldh1a1*, and *Aldh1a3* across all sST datasets. These genes were selected as they serve as markers for specific cell types, such as melanocytes, lens, presumptive neural retina (pNR)2, and pNR3 (Figure 4c, Supplementary Figure 15). Although the absolute position for some of the region is not exactly the same but their relative position remains consistent, such as *Aldh1a3* located in rostral ragion of the retina layer while *Aldh1a1* patterned towards caudal region (Supplementary Figure 16).

In addition, we obtained snRNA-seq data with eye region sectioned as input. However, a limited number of cells were found to express the aforementioned genes, and these cells were primarily clustered in the lower corner of the UMAP plot (Figure 4d, Supplementary Figure 15e). Unfortunately, we were unable to further categorize this small subset into more detailed subgroups, as was the case with the sST data. Interestingly, we also noted that *Crybb3*’s expression values in the snRNA-seq data were relatively lower than expected. This is in line with our earlier observation that *Crybb3* was not found to be expressed in the eye area captured by Visium, indicating a potential capture bias associated with 10X technology.

While snRNA-seq data may not capture as many cells in the eye area as sST methods do, it serves as a useful reference dataset for annotating the sST data. As illustrated in Supplementary Figure 17, the integration of snRNA-seq data with sST data using *Seurat* aided the annotation of sST data. For instance, it improved the annotation of epithelium cells in Stereo-seq data, which had been relatively challenging due to an unknown cluster of cells with mixed expression profiles. This cluster was better resolved using the projection of snRNA-seq epithelium cells (Supplementary Figure 17a). Additionally, the projection facilitated the separation of melanocytes and epithelium cells in BMKMANU S1000 data (Supplementary Figure 17b).

Another issue that deserves mention is the susceptibility of sST technologies to blood contamination, which is often introduced during the tissue preparation and sectioning process and is difficult to avoid. In contrast, snRNA-seq can mitigate this effect using microfluidic techniques. We used the *Hba-a1* gene as an example to evaluate the influence of blood contamination in these sST methods. Our findings revealed that Visium, followed by BMKMANU S1000, were significantly impacted by blood contamination, with all Visium spots and 70% of BMKMANU S1000 spots expressing *Hba-a1*. In contrast, Stereo-seq data exhibited a relatively similar level of blood contamination compared to snRNA-seq, and Slide-seq V2 had the lowest amount of blood contamination (Supplementary Figure 18).

### 2.5 Marker gene detection across technologies

Prior studies have underlined the effectiveness and robustness of using a Wilcoxon rank-sum test when identifying marker genes [32]. We employed this test within *Seurat* to find marker genes between clusters. Analysis of top marker genes reveals technology-specific biases in the selection of these markers. For instance, *Pax6*, a transcription factor known as a master regulator of neural lineages, particularly in the retina [33], exhibited variations in representation among different technologies. Specifically, Stereo-seq data highlighted *Pax6* exclusively in the pNR3 cluster, whereas Slide-seq V2 and BMKMANU S1000 data depicted *Pax6* expression across the entire neural retina (pNR1-4) (Supplementary Table 3), consistent with existing literature. This observation underscores the influence of technology choice on the identification of top markers for specific cell types or clusters. Similarly, disparities were observed in the expression of Hes genes in progenitors of the neural retina and *Sox2* in pNR1 and pNR2 (Supplementary Table 3).

The analysis of clustering results on downsampled data has shown that general cell subsets can still be adequately retained even with fewer sequencing reads. However, it appears that a few subsets, particularly those sharing similar expression profiles are a challenge to be clearly separated. To further investigate the effects of downsampling, we compared the marker genes identified in downsampled data with those in the full dataset. We selected 2 pairs of cell subsets to compare the detection performance for cell subsets with relatively similar expression profiles and those that are more distinct. We employed conducted the marker gene detection in two scenarios: 1) pNR4 and pNR1, which exhibit higher similarity, and 2) lens and melanocytes, which have lower similarity, to identify marker genes (Figure 4e).

Our observations revealed that the number of marker genes increased as the number of reads increased in both pairs of comparisons. Notably, the increase in marker genes was more pronounced with deeper sequencing, as illustrated in Figure 4f and Supplementary Figure 19. The ranking of marker gene detection performance across sST methods aligns with the results depicted in Figure 2d in the comparison between cell subsets with relatively distinct expression profiles. In particular, Slide-seq V2 exhibits higher sensitivity (Figure 4f, bottom panel). Furthermore, our analysis identified a set of genes consistently identified as marker genes across different platforms in each of the comparison pairs. However, each platform exhibited a great number of unique marker genes as well (Figure 4g).

Cell-to-cell communication was applied afterward, but no consistent results could be found across the communication methods applied including *CellChat* [34] and *CellPhoneDB v4* [35] and sST methods (Supplementary Figure 20).

## 3 Discussion

Evaluating spatial transcriptomic methods is more challenging than evaluating scRNA-seq methods. First, it is harder to design a reference tissue for spatial transcriptomics. For scRNA-seq, one could use cell line mixtures/PBMC samples [8, 36, 37], or even purified and diluted mRNA to obtain consistent inputs for different technologies [36]. For spatial transcriptomics, if we use genuine tissues with clear cell type and gene expression patterns, the position and ground truth are then less obvious and limited by our understanding of reference tissues. Second, the measurements are not performed on the same unit. For methods like Visium, the diameter of a spot is larger than 50 microns resembling a mini-bulk RNA-seq. For methods such as Stereo-seq, the spot size is sub-micron, which is much smaller than a single cell.

We carefully designed our benchmarking study to address these challenges. For the first problem, we selected a set of reference tissues with the following criteria: 1) the tissue should be from the widely used model organism, accessible at most research institutes; 2) the tissue should have stable cell-type patterns and specific marker gene expression; and 3) the reference region should have clear morphology that is easy to find in sectioning. Together with the reference tissue, we developed a sectioning protocol to help people reproduce and generate comparable data in the future. For the second challenge, we employed multiple benchmarking metrics and workflows to compare different methods on the same tissue region. We used both all-reads and downsampled data in our comparisons. Downsampling was implemented to mitigate the impact of variations in sequencing depth and cost, however as this may not bring all methods to the same standard as the required number of reads to achieve satisfactory results may differ, we also used all reads in the analysis as complementary results.

In this study, we generated *cadasSTre*, a cross-platform dataset for sequencing-based ST benchmarking that allowed systematic evaluation of 6 sST methods across 22 experiments. We compared various aspects of data from basic metrics to downstream analysis, ranging from sensitivity, and diffusion to clusterability and marker gene detection (Figure 5). Our results suggest spatial transcriptomics requires more sequencing to reach saturation and data generated in this study are well below the saturation level. Technologies such as Stereo-seq require much more sequencing cost to generate high-quality data. Stereo-seq shows the best capture efficiency with raw sequencing depth while Slide-seq v2 gives the best capture efficiency with normalized sequencing depth. Interestingly, we found unexpected gene capturing bias on the Visium platform, with marker genes consistently captured by other technologies not showing up in the Visium data. Considering Visium is the most widely used commercial platform, it is important to further verify its gene-capturing bias on other tissues.

The spot size has become an important metric as a surrogate of the resolution for each method. However, in this study, we highlighted diffusion as a key factor that affects the actual resolution. We found different technologies show distinct diffusion profiles on different tissues. For example, Stereo-seq gives excellent diffusion control on mouse embryo tissue but has much stronger diffusion-induced artifacts in mouse brains. Permeabilization time has a great impact on the molecule diffusion of samples and the tissue-to-tissue variations in diffusion could be a result of it. Although some technologies have sub-micron spot sizes, their real resolution would never reach the same level due to limited sensitivity and high diffusion. Further development of sST would benefit from increased diffusion control and improved assay to determine the permeabilization condition and time, which is a key factors in sST technology development.

Overall, our study generated the first systematic benchmarking scheme of sST methods. Although we strive to make the most of this study, there remain several areas that could be further improved. Also, our understanding of mouse eye development is still limited, making it hard to construct a ground truth in mouse embryo data. The benchmarking dataset generated in this study could be used to further compare computational tools, but it is important to use diverse tissue and data to develop more generalized spatial tools. Although the goal of this study is not to comprehensively benchmark computational tools, we found that clustering tools designed for spatial data may not give better performance than clustering methods for single cells, which agrees with a comparison study [31]. We also found that cell annotations derived from single-cell references may not yield detailed cell states and that clustering derived from spatial data could give complementary results that were sometimes better at resolving rare cell states with spatial patterns. It is important to consider both analyses with and without single-cell references in annotating spatial data.

The sST field is rapidly evolving and the performance of each technology is likely to change with time as they are further optimized. Continuing evaluation is required to keep pace with this fast-moving field. Spatial multi-omics methods are still in their early stages of development [38–41], and new technologies need to be established. Therefore, we believe a community-driven spatial benchmarking league would be beneficial to the future of spatial multi-omics. Our benchmarking efforts highlight the benefits and current issues in the sST field, set up standards for comparing sST methods, and take the first step towards benchmarking spatial multi-omics technology.

## 4 Methods

### 4.1 Sample preparation

#### 4.1.1 Reference sample

All relevant procedures involving animal experiments presented in this study are compliant with ethical regulations regarding animal research and were conducted under the approval of the Animal Care and Use Committee of Westlake University (license number AP#23-111-LXD). Animals were group housed with a 12-hour light-dark schedule and allowed to acclimate to their housing environment for two weeks post arrival. Mouse embryos were collected from pregnant C57BL/6J female mice at embryonic day 12.5 (E12.5). Mouse brain was dissected from 8-week-old C57BL/6J male mice.

#### 4.1.2 Sample preparation, embedding, sectioning, and histological testing

##### Mouse embryo

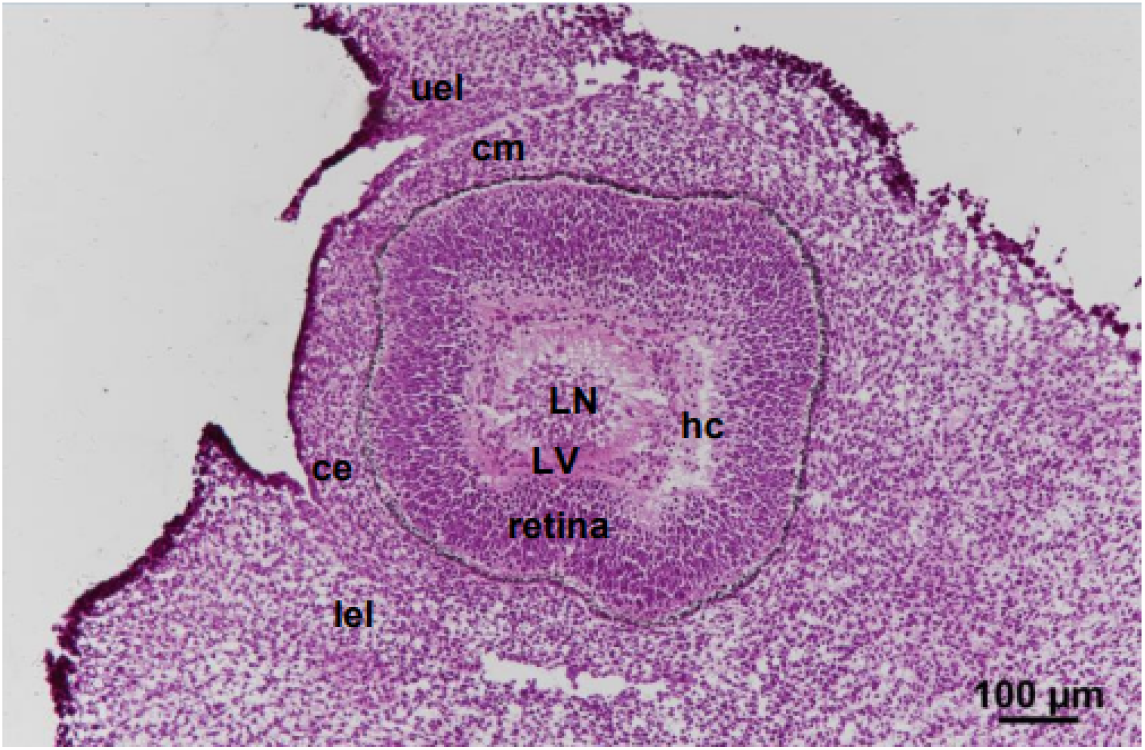

1) E12.5 pregnant female mice were anesthetized with carbon dioxide, and the whole uterus was collected and washed 3 times in ice-cold DPBS; 2) The uterus was separated under a stereo microscope, and each embryo was numbered and photographed with a Motorized Fluorescence Stereo Zoom microscope (ZEISS, Axio Zoom V16); 3) A yolk sac was collected to extract DNA for genotyping (identification of sex); 4) Using dust-free paper to gently wipe the liquid on the surface of the embryo, the embryo was rinsed with ice-cold Tissue-Tek OCT (Sakura, 4583), and then moved to the encapsulation box with ice-cold OCT; 5) Air bubbles were carefully removed with the syringe, and the embryo was placed in sagittal position with tweezers; 6) The location of the embryonic eye was circled and marked the orientation of the embryo, then tissues were transferred to a -80°C freezer, snap-frozen, and stored; 7) Embryos of average size and normal phenotype were selected for subsequent cryosectioning and sequencing (note: the embryos used in our benchmarking analysis came from a litter of mice); 8) Before sectioning, the tissue block was removed from the -80°C freezer and placed in a cryostat (Leica, CM1950) to balance for at least 30 min; 9) The tissue block was smoothly glued to the sample head so that the embryo was sectioned in a sagittal position. If necessary, the angle can be fine-tuned so that the blade section is strictly parallel to the cross-section of the tissue block; 10) Cryosections were cut at a thickness of 10 *µ*m, both the left eye and right eye can be collected; 11) The structure of the sequenced cryosections is shown in the following image: 12) H&E staining procedure: cryosections were balanced at room temperature for 30 min, and then fixed with 4% PFA for 3 min. Then, the sections were washed with ddH_2_O for 2 min, stained with hematoxylin for 6 min, washed with ddH_2_O, stained with eosin for 2 min, washed with ddH_2_O. After that, sections were gradient dehydrated (75% ethyl alcohol for 1 s, 85% ethyl alcohol for 1 s, 95% ethyl alcohol for 1 s, 100% ethyl alcohol for 1 s, 100% ethyl alcohol for 1 min), cleared (xylene for twice), and sealed with Permount TM Mounting Medium after airing. Finally, the figure was scanned using a Motorized Fluorescence Microscope (Nikon, Ni-E).

##### Mouse brain

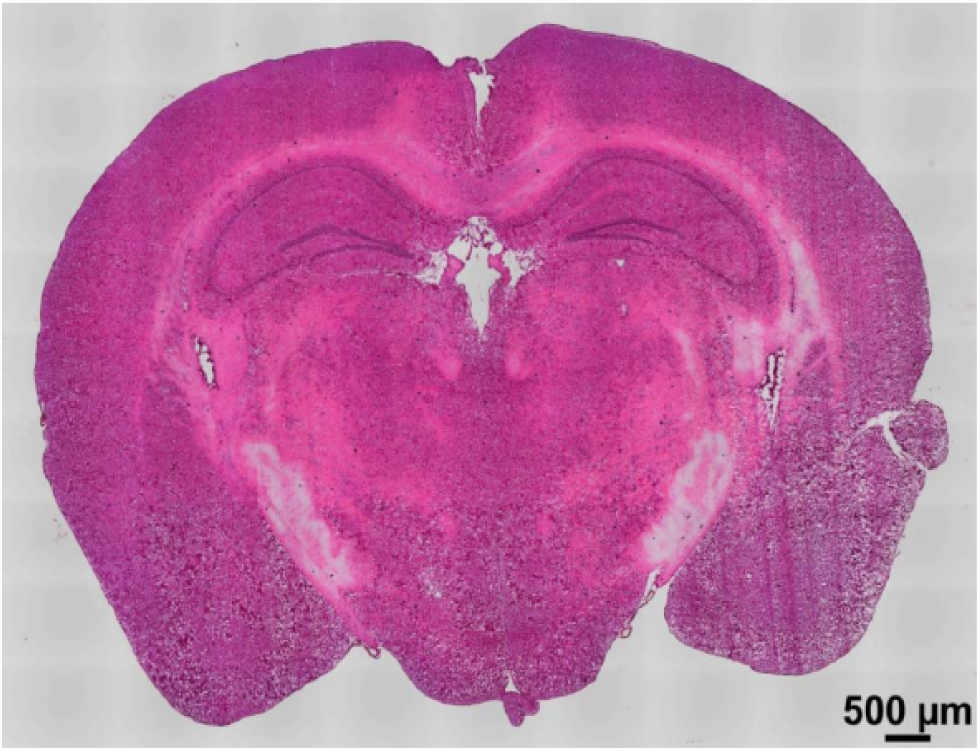

1) 8-week-old male mice were anesthetized with carbon dioxide and decapitated; 2) The whole brain was rapidly dissected, numbered, and photographed with a Motorized Fluorescence Stereo Zoom microscope; 3) Using dust-free paper to gently wipe the liquid on the surface of the brain, the brain was rinsed with ice-cold Tissue-Tek OCT (Sakura, 4583), and then moved to an encapsulation box with ice-cold OCT; 4) Air bubbles were carefully removed with the syringe, and the brain was placed properly with tweezers; 5) The location of the hippocampus was circled and marked the orientation of the brain, then tissues were transferred to a -80°C freezer for snap-frozen and storage; 6) Brains of average size and normal phenotype were selected for subsequent cryosection and sequencing; 7) Before sectioning, the tissue block was taken out from -80°C freezer and placed in a cryostat (Leica, CM1950) to balance for at least 1 h; 8) The tissue block was smoothly glued to the sample head, and the cerebellum was oriented towards the experimenter so that the brain was sectioned in a coronal position. If necessary, the angle can be fine-tuned so that the blade section is strictly parallel to the cross-section of the tissue block; 9) Cryosections were cut at a thickness of 10 *µ*m; 10) The structure of the sequenced cryosections is shown in the following image: 11) H&E staining procedure: cryosections were balanced at room temperature for 30 min, and then fixed with 4% PFA for 3 min. Then, the sections were washed with ddH_2_O for 2 min, stained with hematoxylin for 6 min, washed with ddH_2_O, stained with eosin for 1 min, washed with ddH_2_O. After that, sections were gradient dehydrated (75% ethyl alcohol for 1 s, 85% ethyl alcohol for 1s, 95% ethyl alcohol for 1 s, 100% ethyl alcohol for 1 s, 100% ethyl alcohol for 1 min), cleared (xylene for twice), and sealed after airing. Finally, the figure was scanned using a Motorized Fluorescence Microscope (Nikon, Ni-E).

#### 4.1.3 In-situ imaging with padlock probes

To validate the expression of marker genes, we performed in-situ hybridization and imaging following a simplified version of targeted ExSeq [42]. More specifically. we used 4 fixed barcode regions for distinct fluorescent probes (FAM6, CY3, TXRED, CY5) so we could detect at most 4 genes at the same time without performing muli-round imaging for in-situ sequencing. The tissue was sectioned on Leica CM1950 Cryostats, with 10-micron sections placed on the CITOTEST adhesion microscope slides. The section was then fixed with 4% formalin for 15 minutes at room temperature and washed two times with PBS. Permeabilization of tissue was done with ice-cold 70% EtOH overnight at -20°C. RNase inhibitor (Lucigen) was added at 0.4 U/*µ*l throughout the incubation until the rolling cycle amplification (RCA) was done. The padlock probe was diluted at a final concentration of 5nM per probe, in wash buffers with 2XSSC and 20% formamide. Hybridization was done overnight at 37°C, then washed with the same wash buffer (2XSSC and 20% formamide) 3 times for 15 minutes each, followed by washing with PBS for 15 minutes at 37°C. SplintR ligase (NEB, M0375) was used for probe ligation at 37°C for 2.5 hours. RCA was performed at 30°C overnight using Phi29 enzyme mix (NEB, M0269L). fluorescent probes hybridization was done at 2X SCC and 10% formamide buffer mix, diluting the probes at 100 *µ*M and incubate at 37°C for 1 hour. Imaging was performed with a NIKON A1 confocal microscope with 10X objectives and 2*×*2 image stitching.

#### 4.1.4 Protocols used in different ST methods

See Table1.

### 4.2 Data processing

#### 4.2.1 Preprocessing

We preprocessed fastq files from multiple platforms using their respective preprocessing pipeline (where provided) and updated *scPipe* to allow sample processing with unified functions for data from different sST technologies starting from fastq files.

Mouse GRCm39 was used as a reference for alignment in each of the pipelines for locally generated data.

**Visium** data were processed with spaceranger (v2.1.0), and aligned with STAR 2.7.10b.

**BMKMANU S1000** is a technology developed by BMKGENE (https://www.bmkgene.com/). Similar to HDST, it uses barcoded beads deposited on patterned array. Data were processed with BSTMatrix (v2.3.j), and aligned with STAR 2.7.10b.

**Slide-seqV2** generated bam files of pucks of mouse eyes (Puck 190926 03) and hippocampus (Puck 191204 01 and Puck 200115 08) were downloaded [15].

**Stereo-seq** data were processed with SAW (v6.1).

**DBiT-seq** data underwent initial filtering using a predefined barcode list, and subsequently, fastq file 1 was restructured to adopt the format of spatial barcodes followed by UMIs. The processed data were further analyzed using *scPipe* (v2.0.0) to generate spot-by-gene count matrices.

#### 4.2.2 Selection of region of interest and downsampling

After acquiring count matrices and associated location data for the datasets generated by the aforementioned sST platforms, we aimed to mitigate the impact of variable sequencing depths and costs. To achieve this, we extracted spots located within consensus regions in reference tissues, specifically the hippocampus in the brain and the eye in mouse embryos, for comparative analysis.

Spot selection was guided by histological images (H&E images), feature plots of total counts, and marker genes (*Pmel* for the eye and *Slc17a7* for the brain). These boundaries were meticulously delineated manually.

Subsequently, spots falling within the predefined boundaries for each sample were isolated and used for downsampling. An equal number of reads were chosen within selected spots for both eye and hippocampus data, based on the readID of the selected reads. These selected reads were then isolated from the BAM files generated in the aforementioned pipelines.

The BAM files were further processed to demultiplex based on spatial barcodes and quantified into matrices using UMIs and aligned gene information, using new functions introduced in *scPipe*. In addition to generating count matrices with an equivalent number of reads across platforms, we also processed reads from each platform in specific proportions using *scPipe*.

#### 4.2.3 Sensitivity and diffusion of marker genes

For each sample, we calculated the sum of total counts within selected regions using both the full set of reads and downsampled reads. To assess marker gene sensitivity, we considered specific genes known to be expressed in the dorsal anterior (DA) region of the hippocampus in adult mice (*Prdm8, Prox1*, and *Slc17a7*), as well as genes known to be expressed in the lens of the eyes (*Vit* and *Crybb3*) and a subset of neural retina cells in the eyes of E12.5 mice (*Aldh1a1*). In each sample, we selected five regions measuring 50*µ*m by 50*µ*m in the eyes and five regions measuring 100*µ*m by 100*µ*m in the hippocampus, where these genes were known to be expressed. We individually summed the total number of UMIs in these selected regions within downsampled count matrices to ensure the number of reads was consistent across platforms.

We then performed pairwise comparisons of UMI counts for detected genes across platforms. For eye samples, genes expressed in any one of the platforms with total counts above the 99th percentile but below the 10th percentile in any other platforms were selected for heatmap plotting using a log_1_ 0 scale.

To investigate the gene bias observed in Visium, as demonstrated in the aforementioned pairwise comparisons between platforms, we focused on genes meeting a specific criterion: those expressed across all other platforms with total counts exceeding the 90th percentile and 80th percentile, yet exhibiting number of counts below 1 with Visium. Our exploration of these genes encompassed an analysis of their attributes, including GC content percentage and gene length, using ANOVA analysis. Additionally, we examined the biotypes of these biased genes.

To assess the spatial distribution of marker genes known to be expressed in specific regions of the reference tissues, we used *Pmel* for eye data, *Ptgds* for brain data, and *Slc17a7* for OB data. This analysis used count matrices generated from all reads. We selected regions with expression of these genes roughly in the middle of the chosen regions (6 modalities of 300*µ*m by 50*µ*m in the OB, 6 modalities of 500*µ*m by 50*µ*m in the brain, and 3 modalities of 300*µ*m by 50*µ*m in the eyes). We summed the expression of the aforementioned marker genes for every 10*µ*m along 50*µ*m within 300*µ*m in OB, 500*µ*m in the brain, and 300*µ*m in the eyes. The UMI counts of these marker genes were then aligned based on the location of peak expression and averaged. Modalities with insufficient counts for the selected marker genes were filtered out before plotting.

After the computation of averaged summed values across modalities, these values were subsequently normalized for each platform and depicted in a density plot with the area under the curve standardized to 1. Subsequently, the left half-width half maximum (LWHM) of the profile was computed within each modality, using nonnormalized expression values across platforms. It is worth noting that we employed LWHM as an evaluation metric, drawing inspiration from the full-width half maximum (FWHM) method [19]. In the chosen region, only the left half-width half maximum was used, as there could be an expression of selected genes, such as *Slc17a7* in OB, on the right side of the section that is biologically expected but not caused by lateral diffusion. Modalities that could not be computed with LWHM were excluded from the plotting process. For diffusion analysis, it is important to note that the DBiT-seq data pertained to E10 embryonic eyes, whereas the other datasets were associated with E12.5 embryonic eyes.

To address the significant diffusion in data generated by Stereo-seq in the mouse brain, we conducted diffusion analysis on its downsampled data, consisting of 14% of all the reads.

#### 4.2.4 Cell type annotation

Low-quality spots with total counts below 30% of the first quantile of total counts are filtered out before normalization, which was carried out using the median number of total counts from each platform as the scaling factor. Subsequently, the top 2,000 highly variable genes were identified using the FindVariableFeatures function and used to scale the data through the ScaleData function. A total of 20 principal components (PCs) were then calculated using RunPCA. To categorize spots in each eye sample, we employed 3 distinct methods, including *Seurat* (v4.3.0), *DR*.*SC* (v3.3), and *PRECAST* (v1.6.2).

**Seurat** initially identified neighbors based on 20 PCs, with a *k*-value of 5 chosen for the *k*-nearest neighbor algorithm in FindNeighbors. FindClusters was subsequently applied with various physical resolutions to group known cell-type spots.

**DR.SC** was applied by setting K (the number of clusters) as 10.

**PRECAST** was applied with the number of clusters specified as 10 and using the SelectModel function to reorganize the fitting results within PRECASTObj.

#### 4.2.5 Integration of sST and scRNA-seq data

We followed the instructions in *Seurat* with parameters reduction =`‘cca’, k.filter = NA, and normalization.method = ‘SCT’ in FindTransferAnchors`. Dims were set as 30 with PCA used as weight.reduction in TransferData.

#### 4.2.6 Evaluation of clustering on downsampled data

The downsampled data were subjected to the same pipeline as described above, leveraging *Seurat* to generate clustering results. These obtained clustering results were subsequently compared to clustering outcomes obtained through the processing of count matrices generated from the entire set of reads. To visualize this comparison, we created a heatmap with a logarithmic scale, illustrating the corresponding number of spots in each of the downsampled clustering and the overall clustering results. The entropy of accuracy and purity were then calculated. *ECA* and *ECP* are defined as follows:

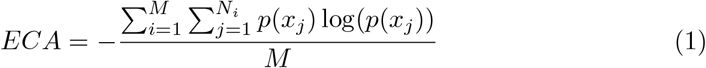

where *M* denotes the number of clusters generated from a method (the clustering solution to be evaluated), *N*_*i*_ denotes the number of elements in the *i*th cluster based on the ground truth (here the provided labels) and

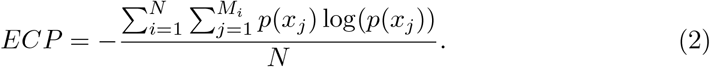

#### 4.2.7 Marker genes detection

FindMarkers in *Seurat* was applied to find marker genes between two pairs of cell subsets: 1) lens and melanocytes; 2) pNR4 and pNR1. Genes that exhibited higher expression in the lens and pNR4, with expression levels exceeding 5% of the specific spots, the log-fold-change greater than 0.25, and an adjusted *p*-value provided by *Seurat* less than 0.01, were considered as marker genes.

#### 4.2.8 Cell communication analysis

Cell communication analysis was then performed on spots with distinct annotated cell types, using methods including *Cellchat* (v1.6.1), *CellPhoneDB* (v4).

**Cellchat** used the CellChatDB database of the mouse, creating cellchat objects based on annotation information, and employed the default ‘Trimean’ statistical method.

**CellPhoneDB** was applied by transforming mouse genes into their human homologs using the biomaRt package. Using the CellPhoneDB database for cellphone analysis, we conducted using 1,000 random permutations in the analysis following the tutorial https://github.com/ventolab/CellphoneDB/blob/master/notebooks/T01_Method2.ipynb. The minimum cell percentage threshold required to consider a gene as expressed in the analysis was set to 0.1, and significance was determined with a *p*-value threshold of less than 0.05.

## Data availability

Raw count matrices are available at the National Genome Data Center (https://www.cncb.ac.cn/) under BioProject accession code PRJCA020621. A summary of individual accession numbers is given in Supplementary Table 2. The *c*adasSTre data collections are continually updated on our website genographix.com. The standard sectioning protocol is deposited in protocols.io: dx.doi.org/10.17504/protocols.io.5qpvo379dv4o/v1.

## Code availability

Scripts used to process the data are available at https://github.com/YOU-k/cadasSTre.

## Acknowledgments

We thank members of the Tian and Liu laboratories for helpful discussions. We thank Professor Yixue Li for his support to the Tian Lab and to this project. We thank Professor Gang Cao for the helpful discussion. This project is supported by Guangzhou National Laboratory (Grant No. YW-YFYJ0301), Westlake Foundation, and the Key research and development project of the Ministry of Science and Technology (2022YFA1105700).

## Author contributions

L.T. and X.L. designed and supervised this study. Y.F. curated reference tissues, wrote sectioning protocol, and performed sST experiments with help from S.J. and X.L. Y.Y. conducted data analyses with help from L.L., Z.Z., Y.X, A.S.K and Y.L. S.L. and W.R. performed padlock probe imaging experiments. F.J. and G.P. performed DBiT-seq experiments. Y.Y., Y.F, M.E.R. and L.T. wrote the manuscript with the contribution of all of the authors.

## Appendix A Supplementary information

Supplementary figures and Supplementary tables.

