## Supplementary Figures for "Systematic comparison of sequencing-based spatial transcriptomic methods"

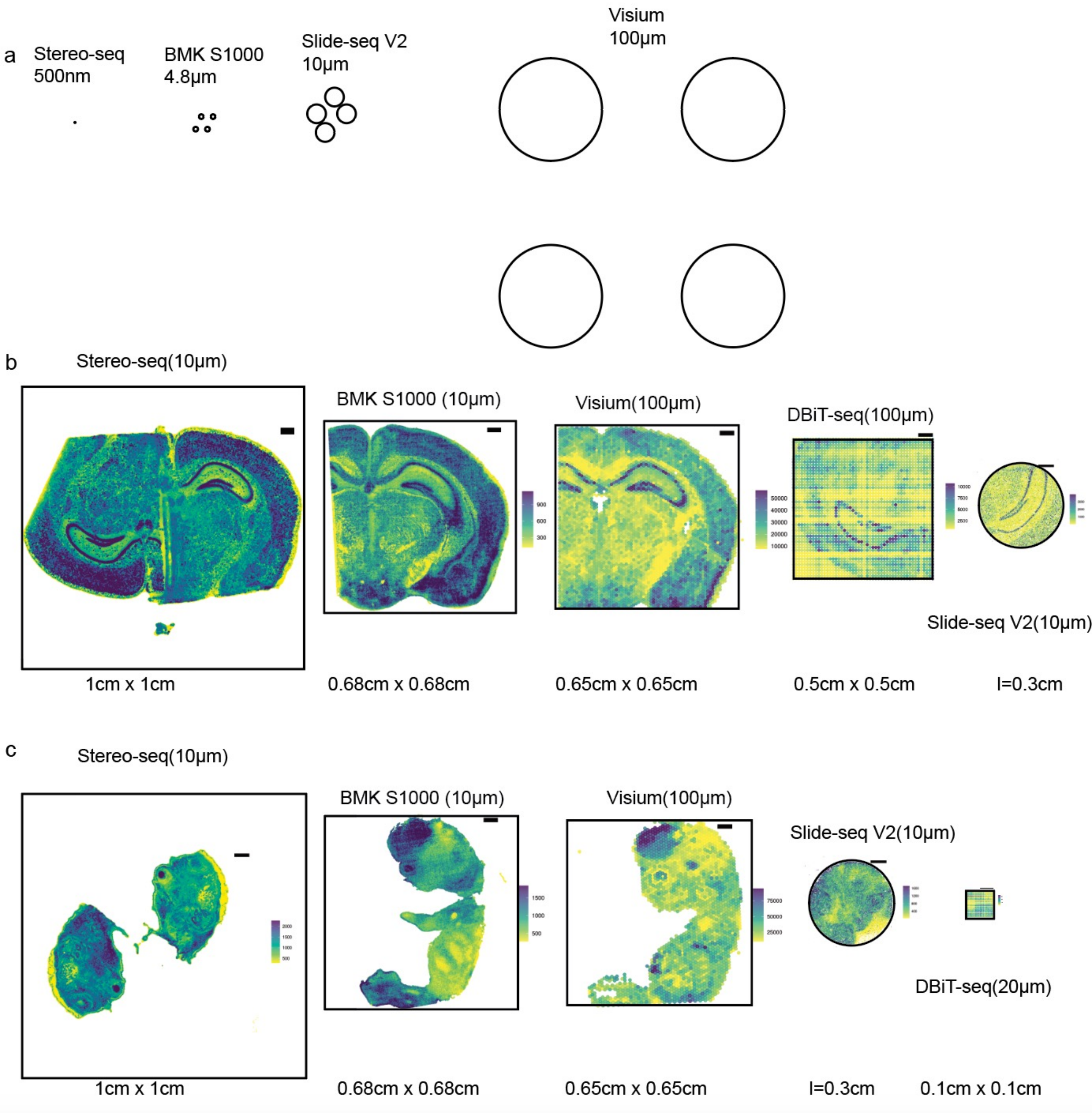


**Supplementary Figure 1**.
a) Spots are plotted according to their respective distances between spot centers and their positions on the chip.
The capture area for each sST method is depicted with a black box, overlaid with b) a brain sample and c) an embryo sample. The size of each box corresponds to the capture area information listed below, offering a comprehensive overview of the differences in capture areas.

### 10X Visium

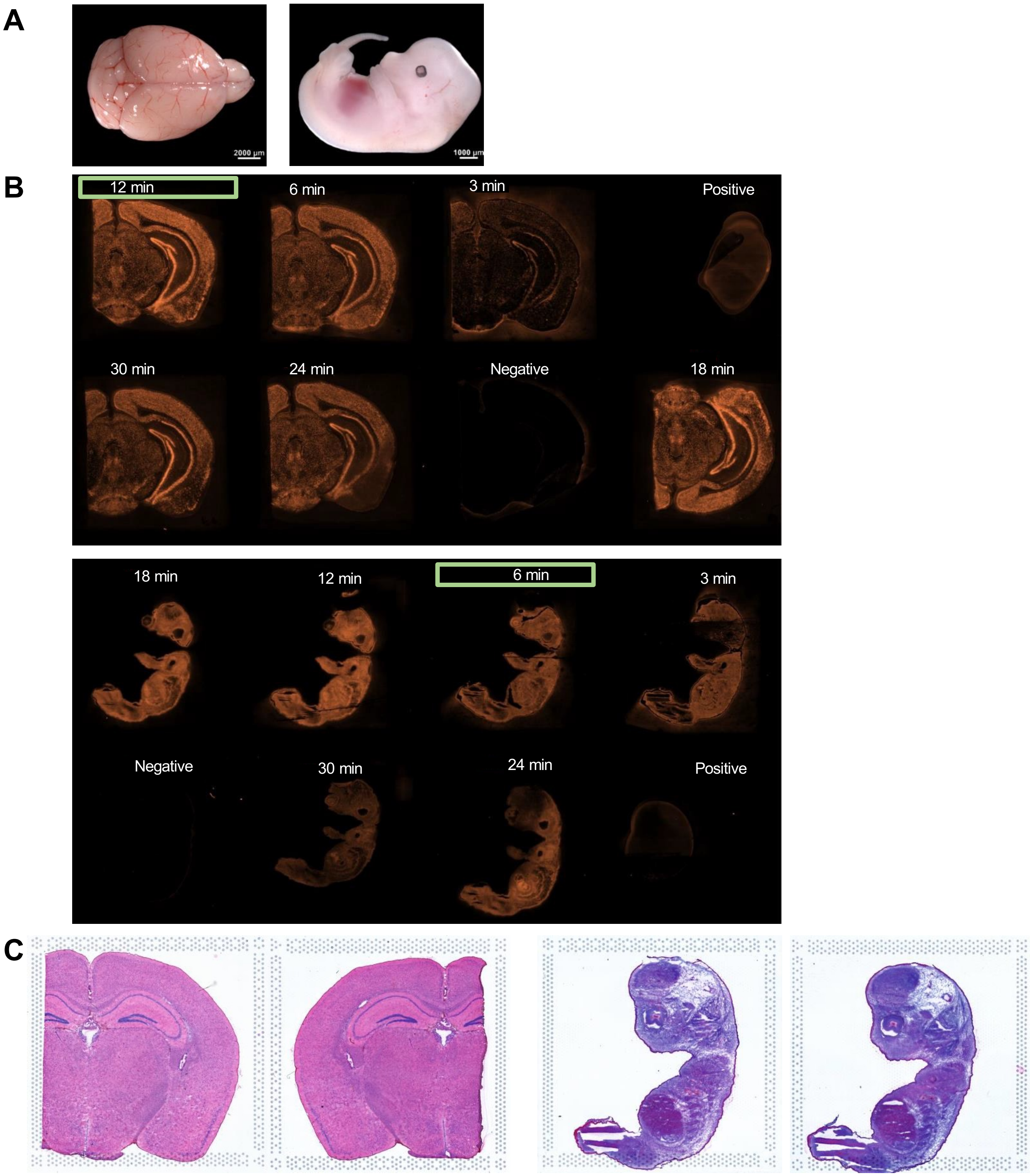


**Supplementary Figure 2**.
A. Tissue blocks used for 10X Visium spatial technologies.
B.Tissue permeabilization were optimized before spatial gene expression experiment. Permeabilization for 3 min, 6 min, 12 min, 18 min, 24 min, and 30 min were performed, and fluorescence value was used for evaluation. 12 min, 6 min permeabilization time was chosen for the spatial gene expression experiment of mouse brain and E12.5 embryonic eye, respectively.
C. H&E staining of sections used for spatial expression.

**Stereo-seq**

**A**

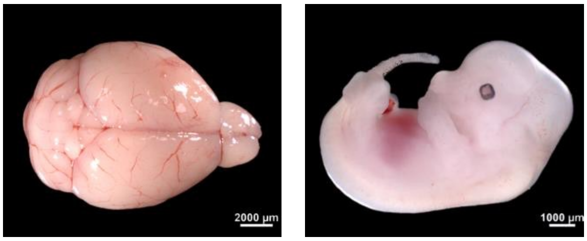


**B**

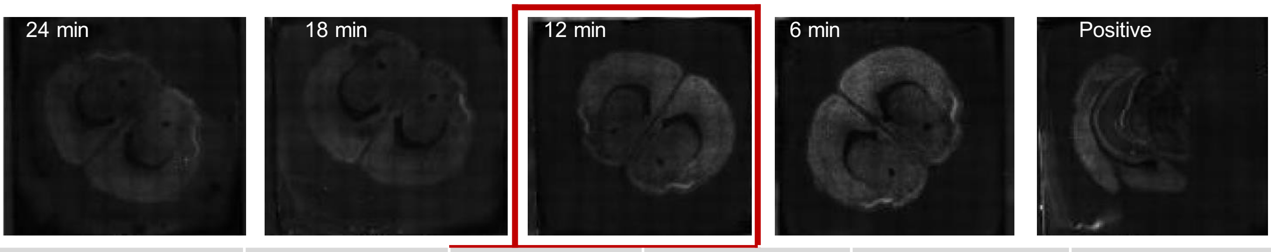


| permeabilization time | 24 min | 18 min | 12 min | 6 min | Positive |
|---|---|---|---|---|---|
| gray value | 22.08 | 25.651 | 25.163 | 28.008 | - |

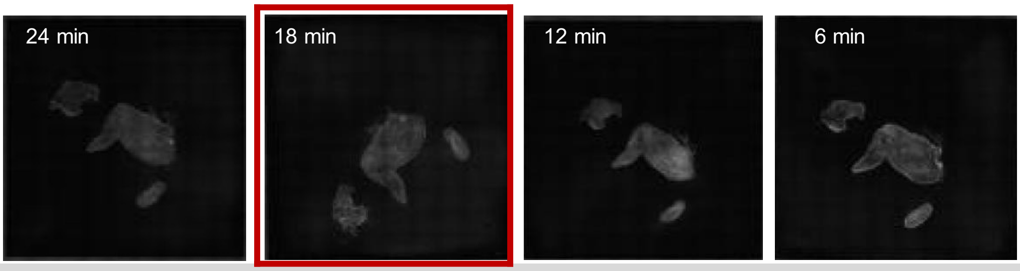


| permeabilization time | 24 min | 18 min | 12 min | 6 min |
|---|---|---|---|---|
| gray value | 13.074 | 18.804 | 12.893 | 15.399 |

**C**

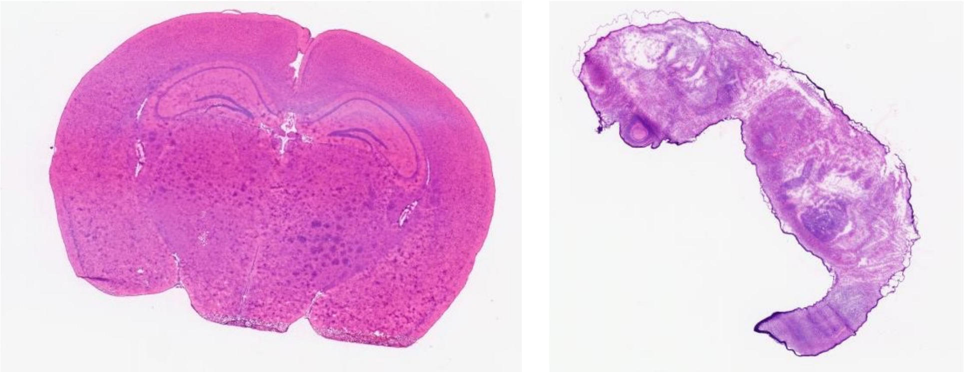

**Supplementary Figure 3**.
A. Tissue blocks used for Stereo-seq spatial technologies.
B.Tissue permeabilization were optimized before spatial gene expression experiment. Permeabilization for 6 min, 12 min, 18 min, and 24 min were performed, and fluorescence value was used for evaluation. 12 min, 18 min permeabilization time was chosen for the spatial gene expression experiment of mouse brain and E12.5 embryonic eye, respectively.
C. H&E staining of adjacent sections used for spatial expression.

BMK S1000

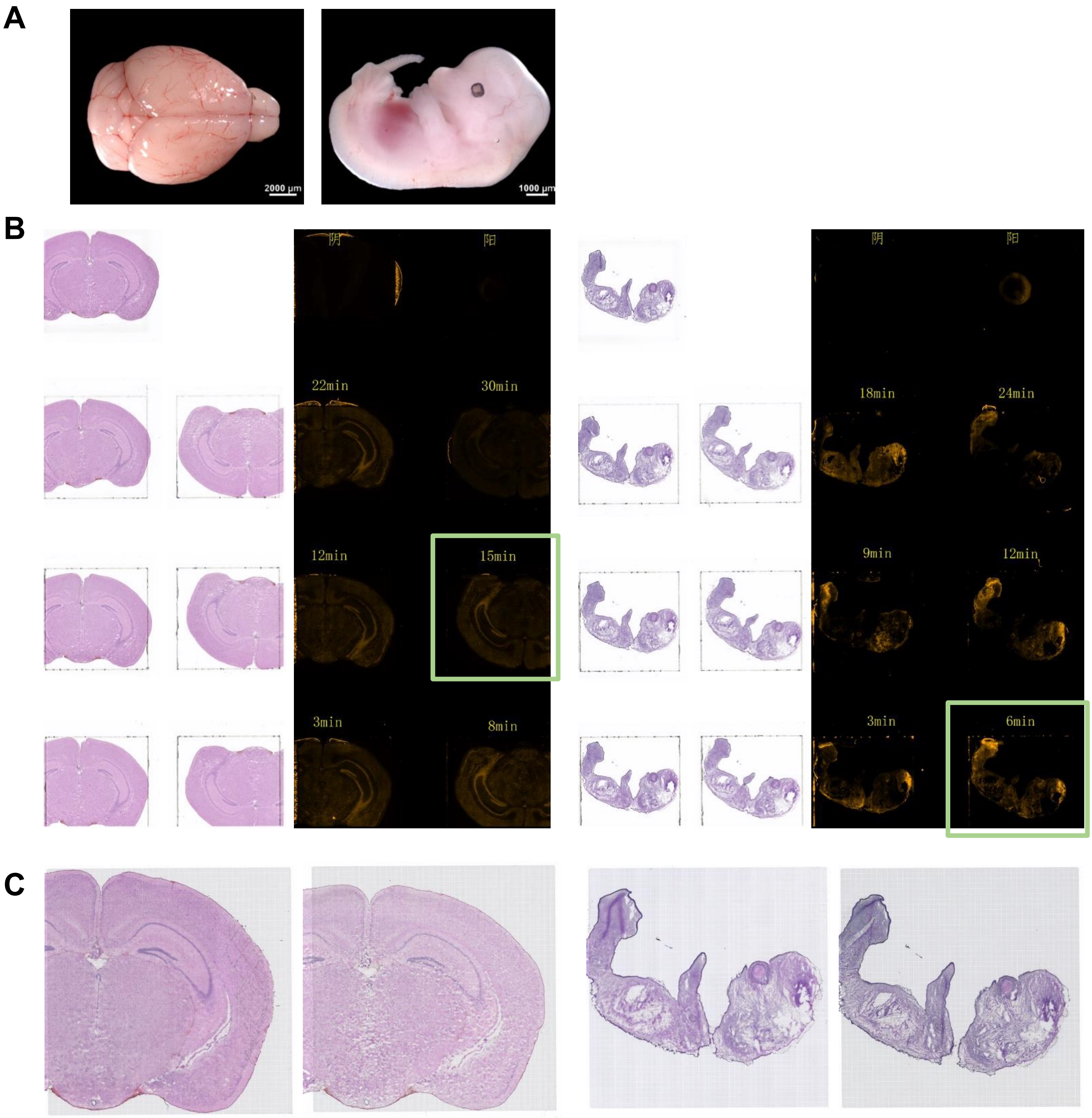


**Supplementary Figure 4**.
A. Tissue blocks used for BMK S1000 spatial technologies.
B.Tissue permeabilization were optimized before spatial gene expression experiment. Permeabilization for 3 min, 8 min, 12 min, 15 min, 22 min, and 30 min were performed, and fluorescence value and corresponding H&E staining was used for evaluation. 15 min, 6 min permeabilization time was chosen for the spatial gene expression experiment of mouse brain and E12.5 embryonic eye, respectively.
C. H&E staining of sections used for spatial expression.

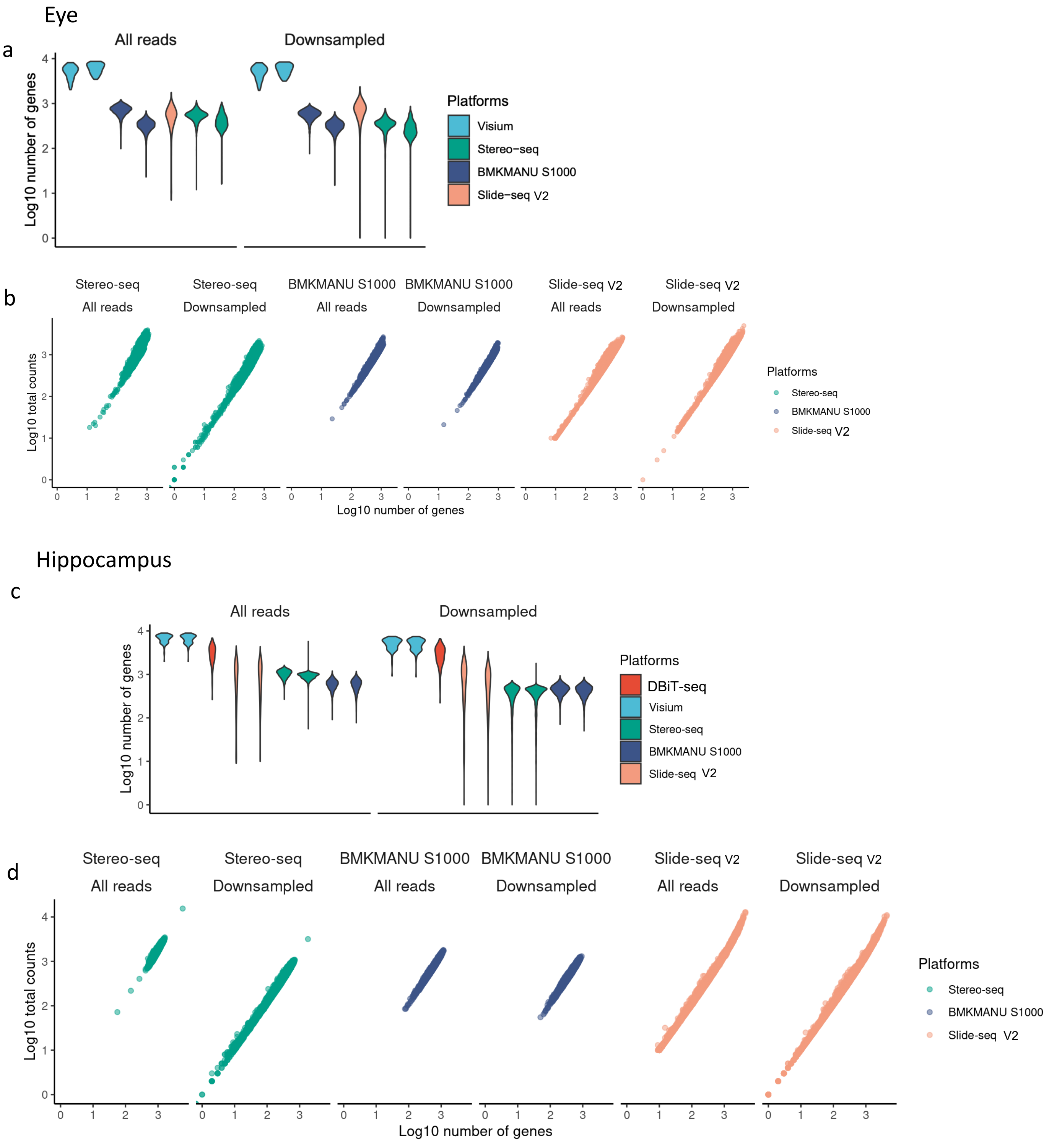


**Supplementary Figure 5**. a) In a selected region of the E12.5 mice eye, the left panel shows the log10-transformed number of detected genes per spot, using data from all reads. In the right panel, the same information is presented using downsampled data. Notably, the spot size for 10X Visium is 50µm x 50µm, while for other platforms, it is 10µm x 10µm. The relationship between the log10-transformed number of genes and log10-transformed total counts per spot is individually plotted in b).

c) For a selected region in the adult mice hippocampus, corresponding plots are provided in c) and d). The same format and spot size considerations apply.

a

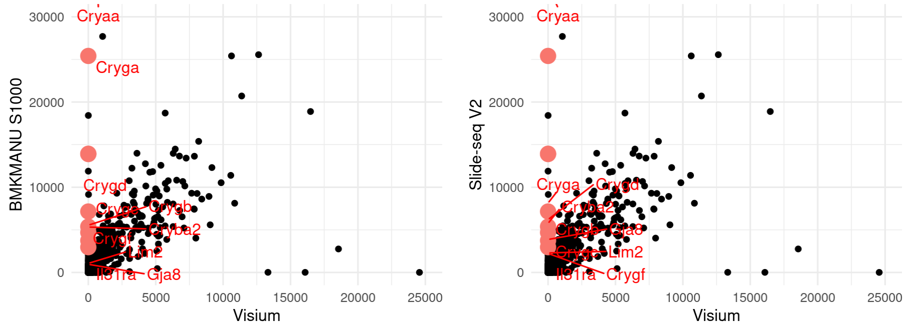


b

*scPipe* processed Visium data

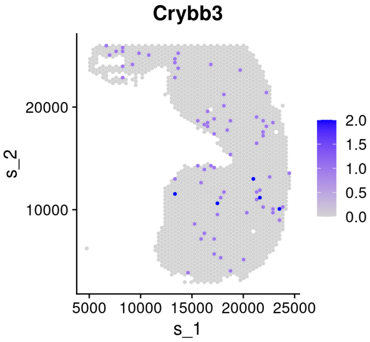


**Supplementary Figure 6**. a) Expression levels of all detected genes are compared between Visium and other platforms. Each dot represents a gene, shown in black. Genes that display expression at the 90th percentile in other platforms but are at the 10th percentile in Visium are highlighted in red and labeled with their gene symbols.
b) Expression of the example gene Crybb3, which exhibited bias as illustrated in plot a), is visualized in spatial reduction. Preprocessing was performed using *scPipe* instead of *SpaceRanger*. Notably, unexpected expression of Crybb3 was observed in Visium data, contrasting with the expected high expression in the lens, consistently observed in data generated by other platforms.

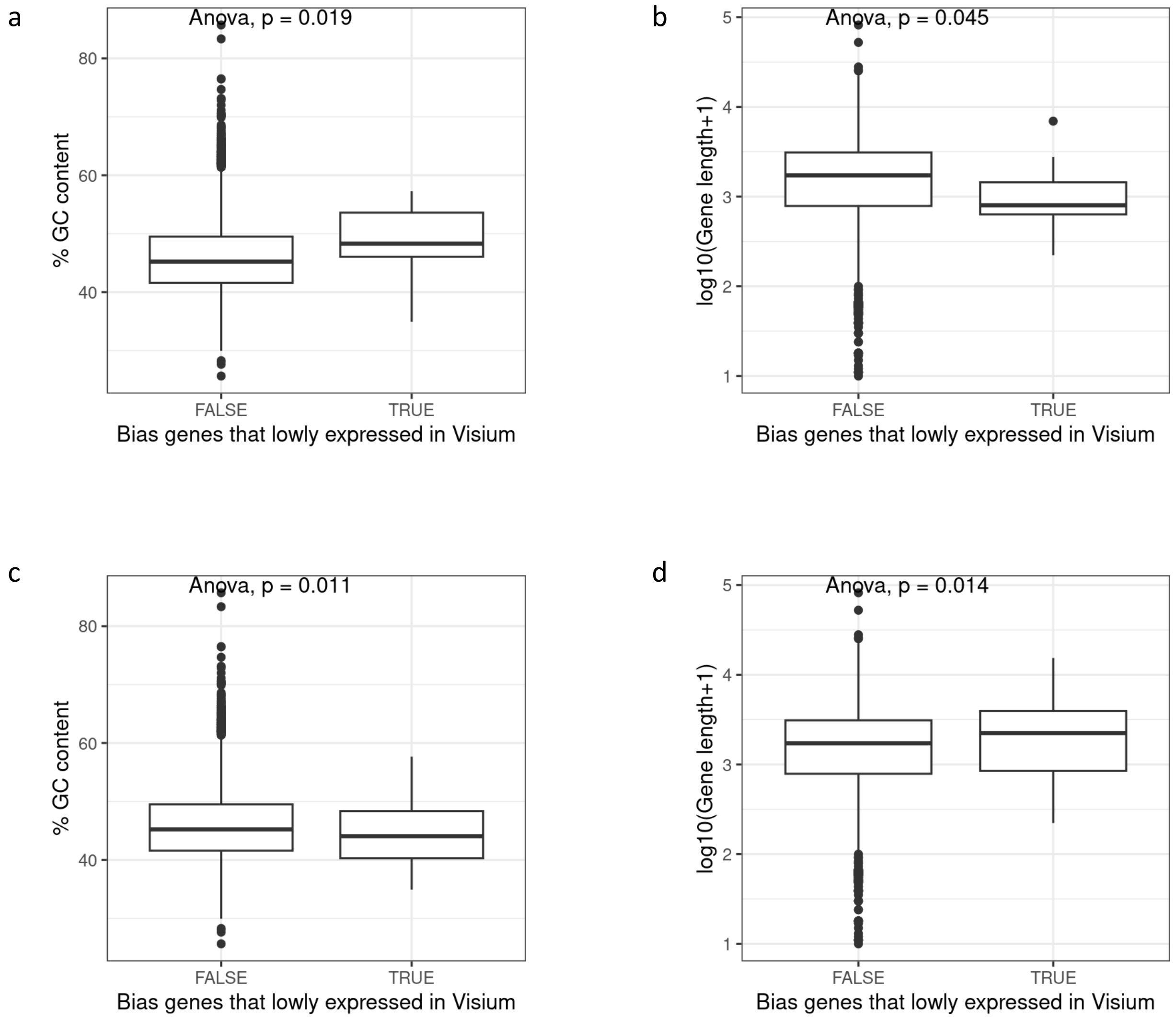


**Supplementary Figure 7**. Genes meeting a specific criterion: those expressed across all other platforms with total counts exceeding the 90th percentile and 80th percentile, yet exhibiting number of counts below 1 with Visium are selected.
ANOVA analysis was performed between GC content percentage and selected genes with a) $90^{th}$ percentile used and c) with $80^{th}$ percentile used .
ANOVA analysis was also performed between gene length and selected genes with b) $90^{th}$ percentile used and d) with $80^{th}$ percentile used.

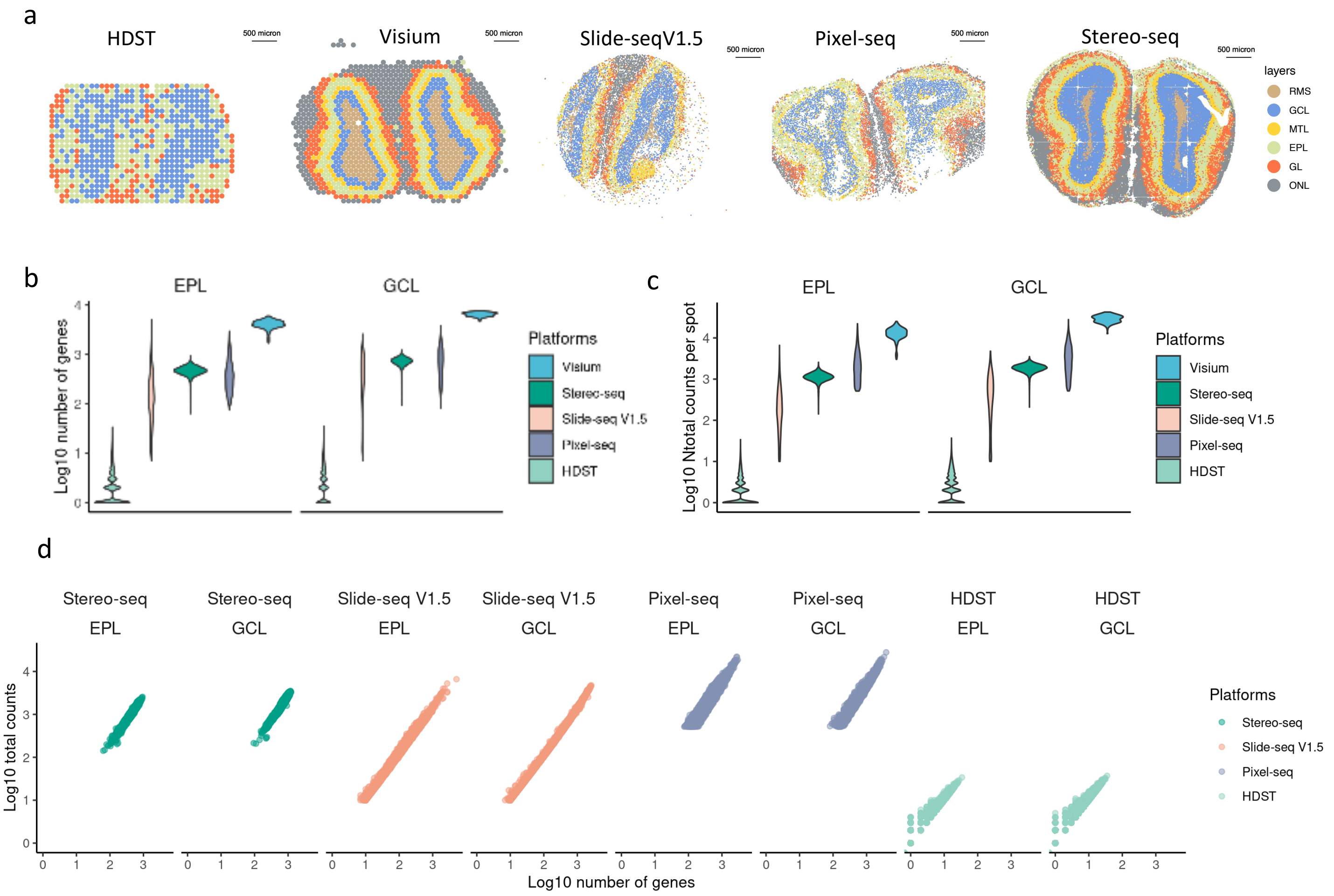


**Supplementary Figure 8**. Data obtained from various platforms for the mouse olfactory bulb were processed and annotated. The spatial reduction plots, color-coded by annotated layers, are presented individually for each platform in a).

In b), the log10-transformed number of detected genes per spot, calculated from the data of all reads used, is depicted separately for the External Plexiform Layer (EPL) and the Granule Cell Layer (GCL). These layers exhibit different gene density and count patterns.

c) Log10-transformed total counts per spot, derived from the data of all reads used, are showed separately for EPL and GCL in c). Notably, the spot size for 10X Visium is 50µm x 50µm, whereas for other platforms, it is 10µm x 10µm.

d) The relationship between the log10-transformed number of genes and the log10-transformed total counts per spot is individually illustrated in b).

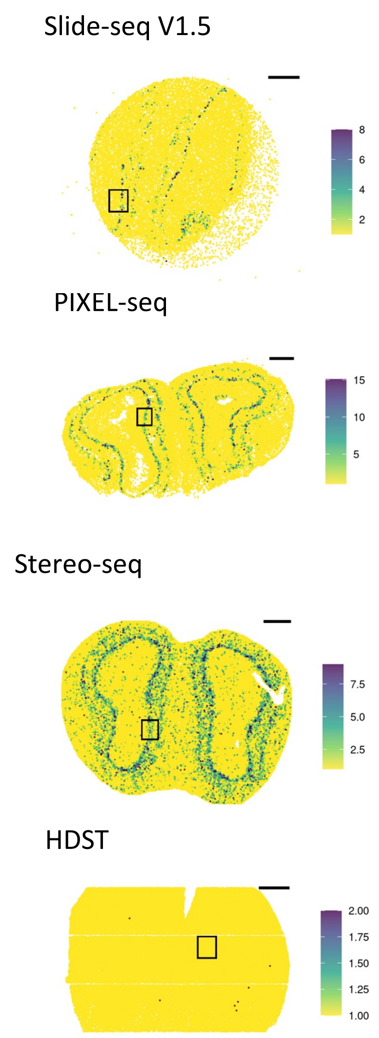


**Supplementary Figure 9**. Spatial reduction plots illustrate the expression of *Slc17a7* across diverse platforms, with colors representing raw count values in mice hippocampus. Black boxes highlight specific regions chosen for diffusion calculations.

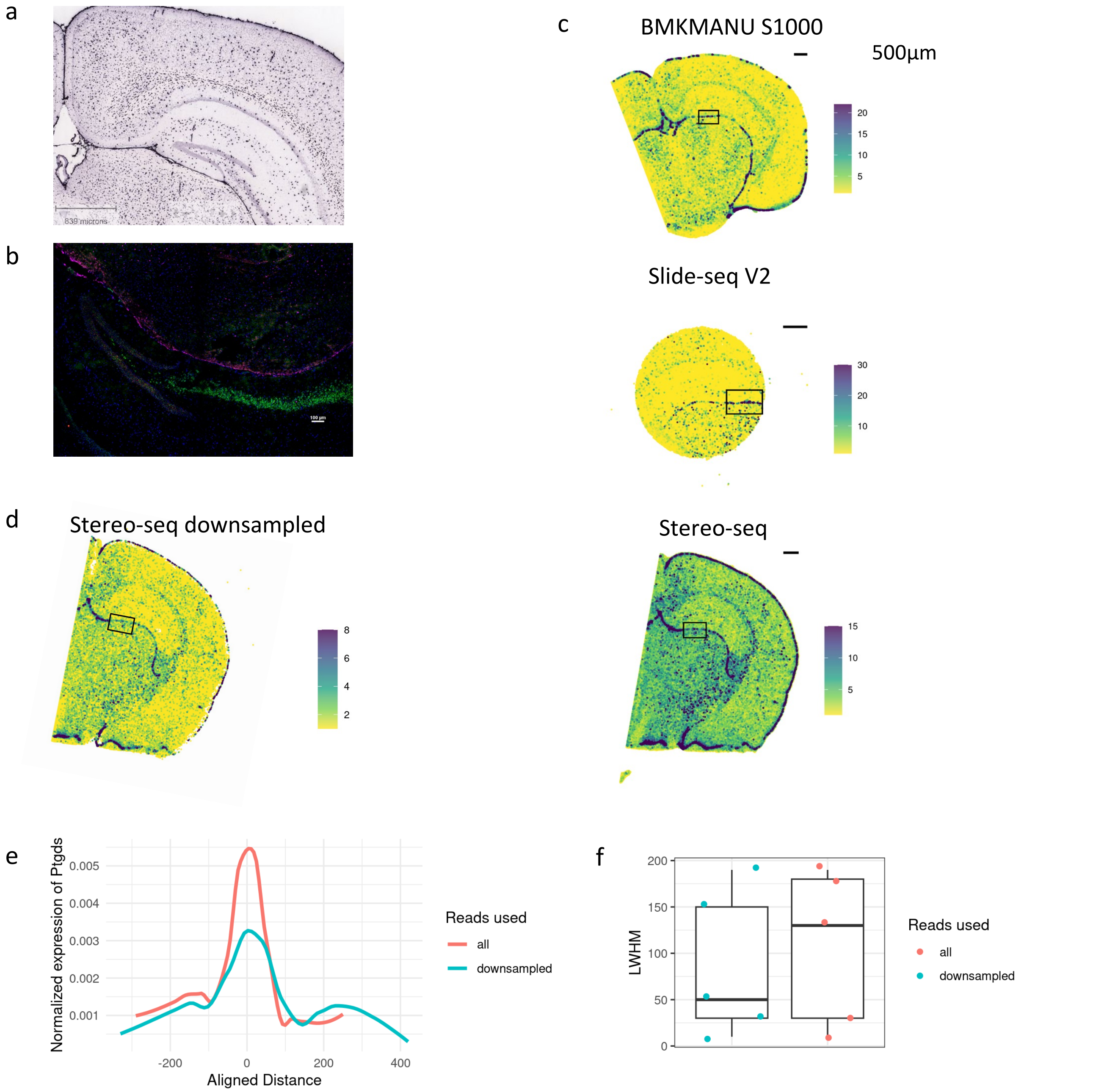


**Supplementary Figure 10**. a) ISH image showing expression of *Ptgds* from [https://mouse.brain-map.org/experiment/show/79567709](https://mouse.brain-map.org/experiment/show/79567709).
b) ISH image with expression *Ptgds* shown in purple, *Prox1* shown in red, *Slc17a7* shown in green and nuclei shown with DAPI.
c) Spatial reduction plots depict the expression of *Ptgds* across different data platforms, with color coding reflecting raw count values. Black boxes delineate specific regions chosen for diffusion calculations.
d) Spatial reduction plots display the expression of *Ptgds* on downsampled data from stereo-seq, generated using reads comprising only 14% of the total number of reads.
e) The average expression of *Ptgds* across selected modalities is illustrated in a density plot. Comparisons are made between results obtained using all reads and downsampled reads.
f) LWHM values were computed for the same sets of modalities and are presented in boxplots, where each dot represents a modality.

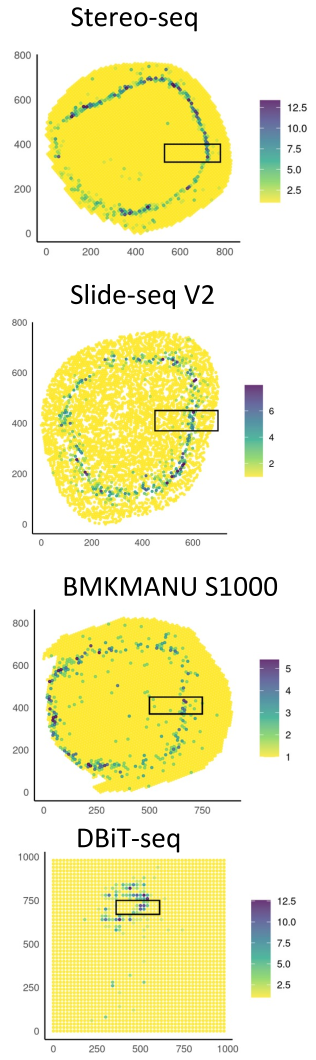


**Supplementary Figure 11**. Spatial reduction plots illustrate the expression of *Pmel* across diverse platforms, with colors representing raw count values in mice embryo eyes. Black boxes highlight specific regions chosen for diffusion calculations.

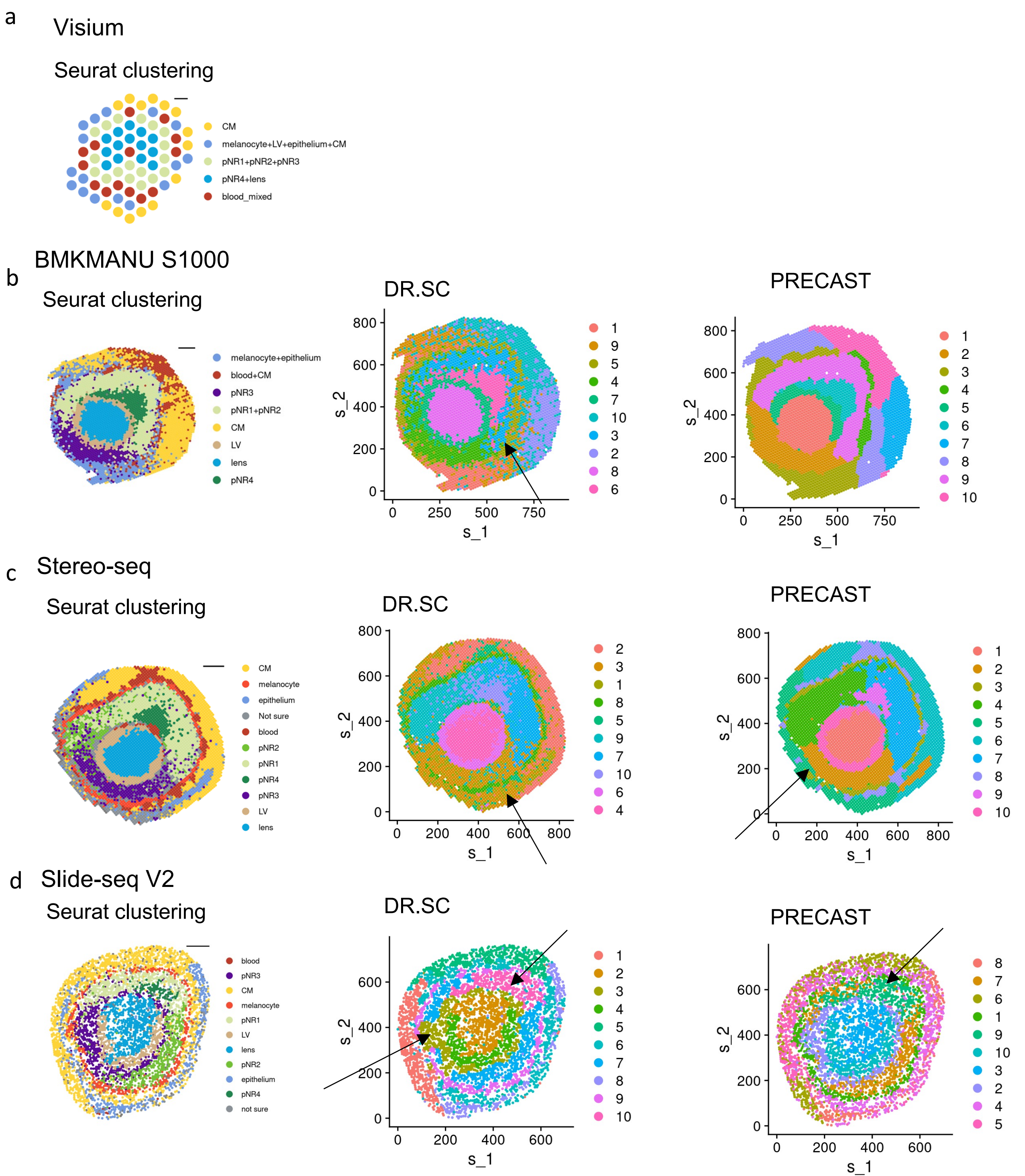


**Supplementary Figure 12**. Count matrices from individual platforms underwent quality control and normalization procedures. Subsequently, clustering was performed using three selected methods. Dimplot visualizations with spatial reduction, colored by clustering results, are presented from left to right, corresponding to Seurat clustering, DR.SC, and PRECAST for the following samples: a) 10X Visium b) BMKMANU S1000 c) Stereo-seq d) Slide-seq V2.

Within each panel, the arrow bar indicates differences between clustering results. Notable findings include:

For BMK data, DR.SC successfully identified a cluster of melanocytes, whereas the other two methods failed to do so.

In the case of stereo-seq data, DR.SC combined pNR3 and epithelium cells and grouped spots with unknown annotation together. PRECAST missed a set of melanocytes located near the region enriched with spots for which annotation remains uncertain.

In Slide-seq V2 data, both DR.SC and PRECAST failed to detect pNR4.

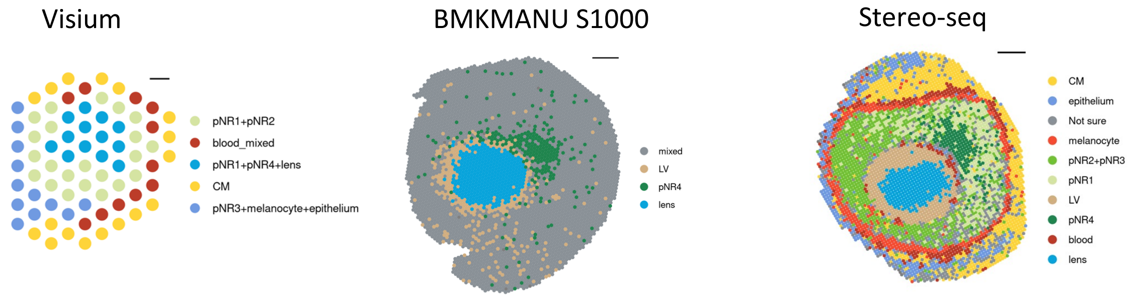


**Supplementary Figure 13**. Expression profiles generated from different platforms are processed and annotated clustering results are shown in their spatial reduction for samples that are not shown in the main plot.

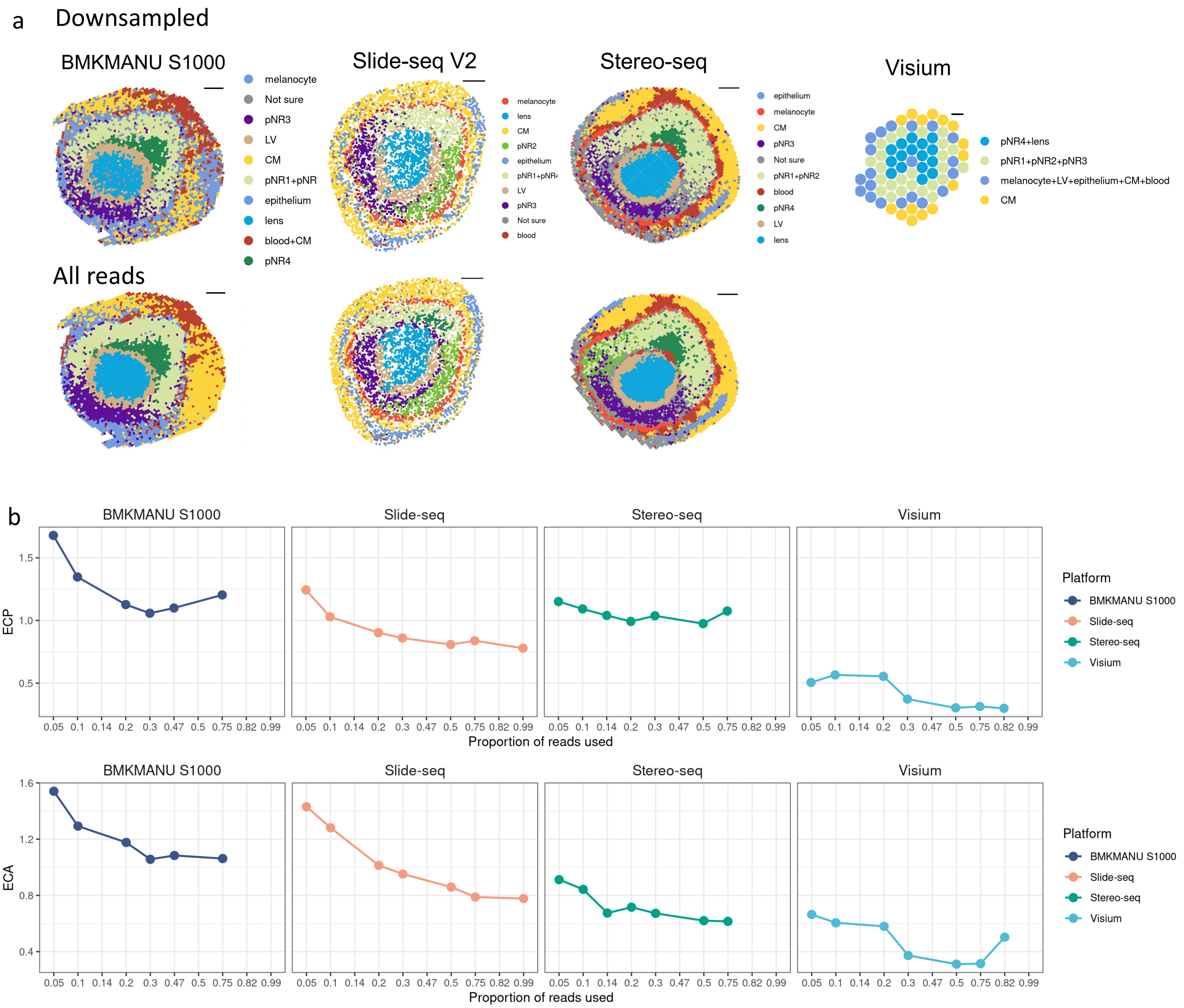


**Supplementary Figure 14**. Downsampled samples, corresponding to those depicted in Figure 3, were utilized for subsequent analyses. Seurat clustering was applied to these downsampled samples, and the clustering results are presented in spatial reduction plots in a), with spots color-coded by annotated cell states. Corresponding spatial reduction plots are displayed underneath in b).

In general, the major cell states are still discernible in downsampled datasets, reflecting the robustness of the analysis.

To assess clustering consistency between downsampled and full-read datasets, ECP (c) and ECA (d) scores were calculated and plotted, with each dot representing a downsampled dataset. Lower ECA and ECP scores indicate more consistent results. Notably, Visium exhibited the most robust clustering results, while data from other platforms displayed greater inconsistency, likely due to the influence of sequencing depth on substates within general cell states.

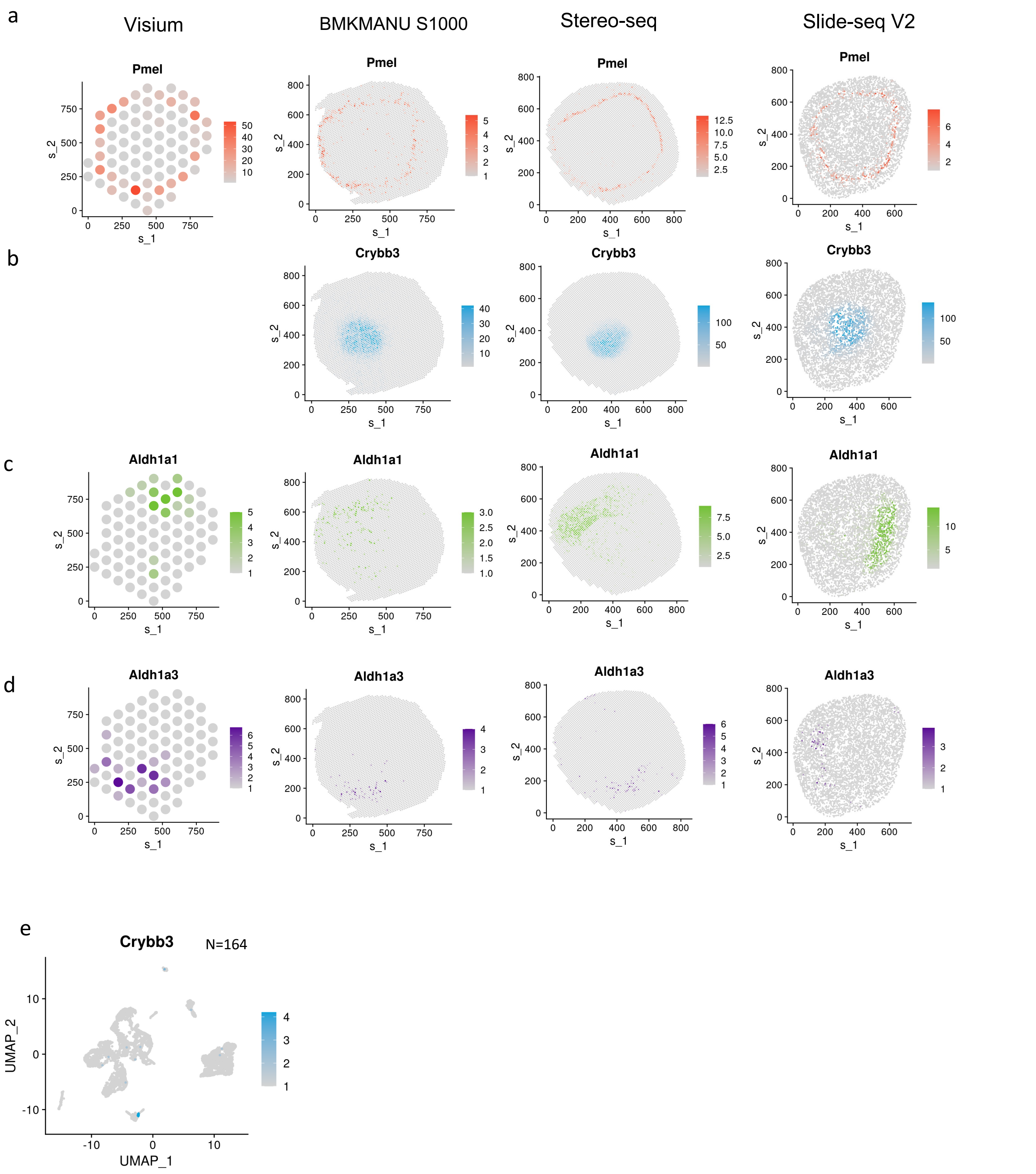


**Supplementary Figure 15**. Expression of *Pmel* a) , *Crybb3* b), *Aldh1a1* c) and *Aldh1a3* d) were plotted in their spatial reduction for data generated from different platforms and the color denotes the values of raw counts of selected genes.

e) Expression plot *Crybb3* was also plotted in UMAP for snRNA-seq data and the number of cells with expression of *Crybb3* above 0 is labelled on the right top.

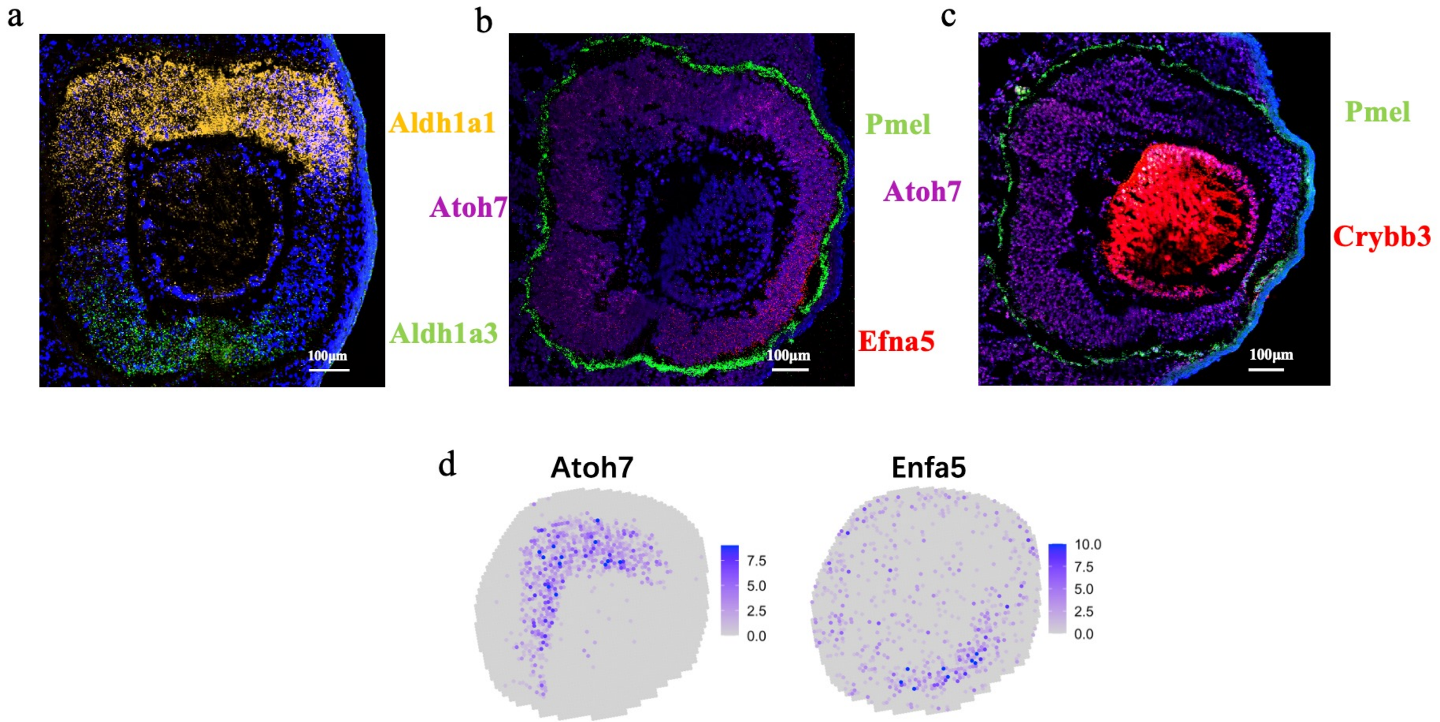


**Supplementary Figure 16**. In situ imaging of E12.5 mouse embryo eye.(a) The orange-red color represents the marker gene Aldh1a1,green represents the marker gene Aldh1a3, blue represents the nucleus. Scale bar, 100 μm. (b) The red color represents the marker gene Efna5, green represents the marker gene Pmel, purple represents the marker gene Atoh7, blue represents the nucleus. Scale bar, 100 μm. (c) The red color represents the marker gene Crybb3, green represents the marker gene Pmel, purple represents the marker gene Atoh7, blue represents the nucleus. Scale bar, 100 μm. (d) Spatial expression of Atoh7 and Enfa5 in Stereo-seq data

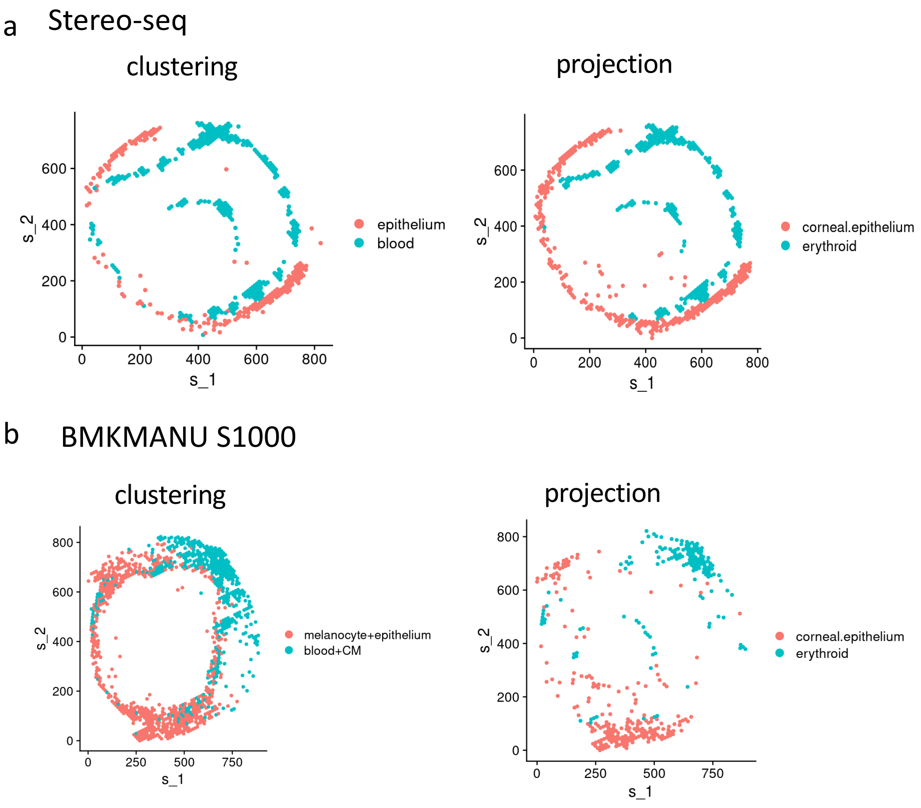


**Supplementary Figure 17**.
Clustering results generated by Seurat related to epithelium and blood and projection results based on snRNA-seq were plotted from left to right for data of a) Stereo-seq, b) BMKMANU S1000 accordingly.

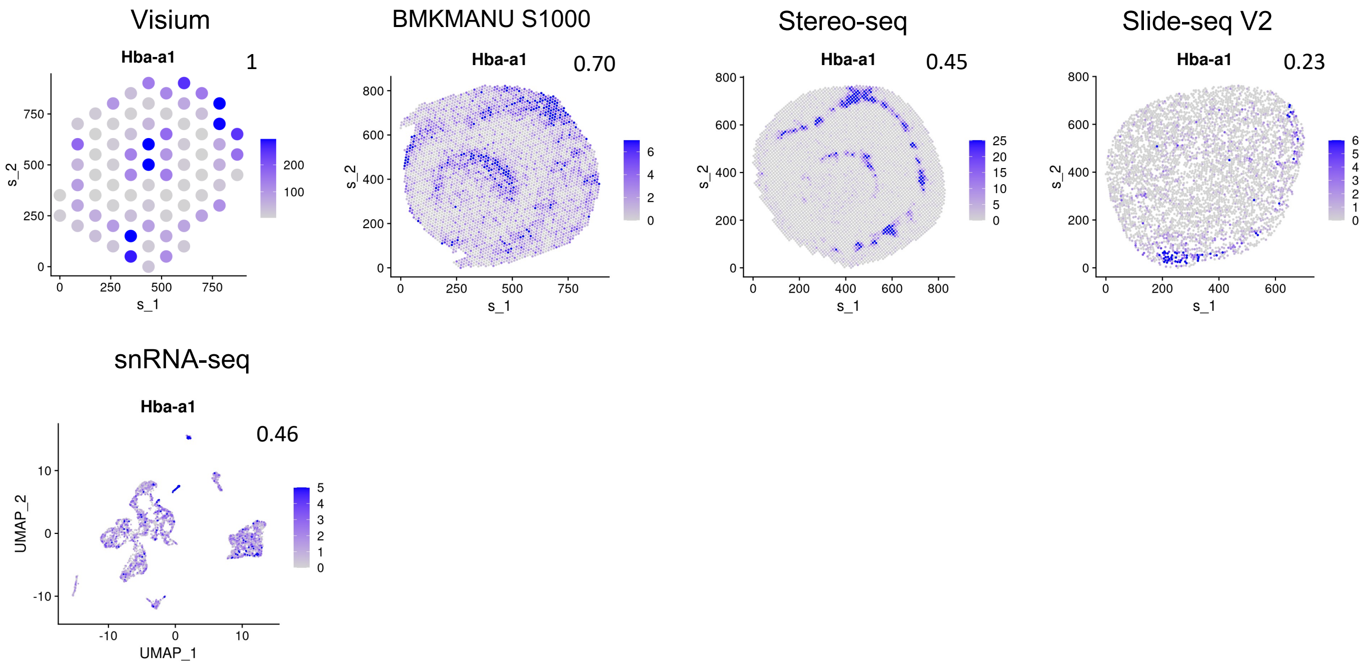


**Supplementary Figure 18** Expression of Hba-a1 in data generated from different plotforms were plotted in their spatial reduction. The color denotes the value of raw expression values of *Hba-a1*. The proportion of cells expressing *Hba-a1* among all cells are labelled on the right-top corner.

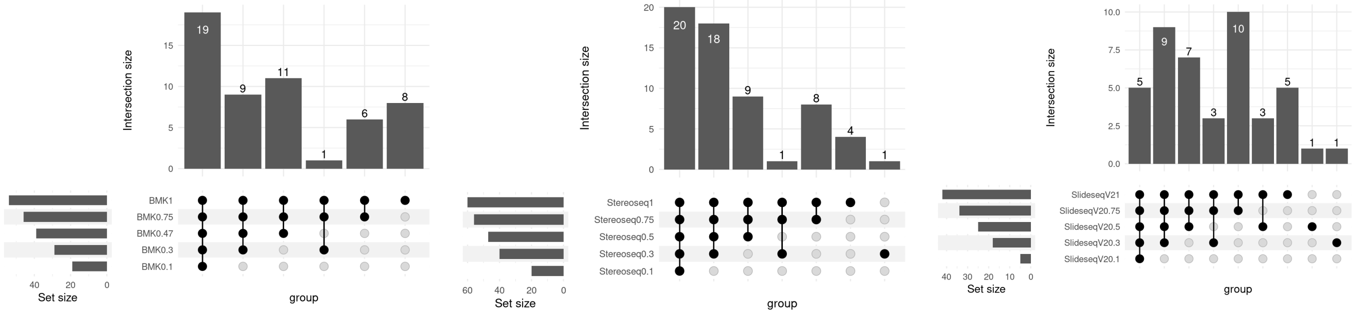


**Supplementary Figure 19**, Upset plots displaying the intersection of marker genes obtained by different sST methods using different proportion of reads for the pNR4 and pNR1 comparison.

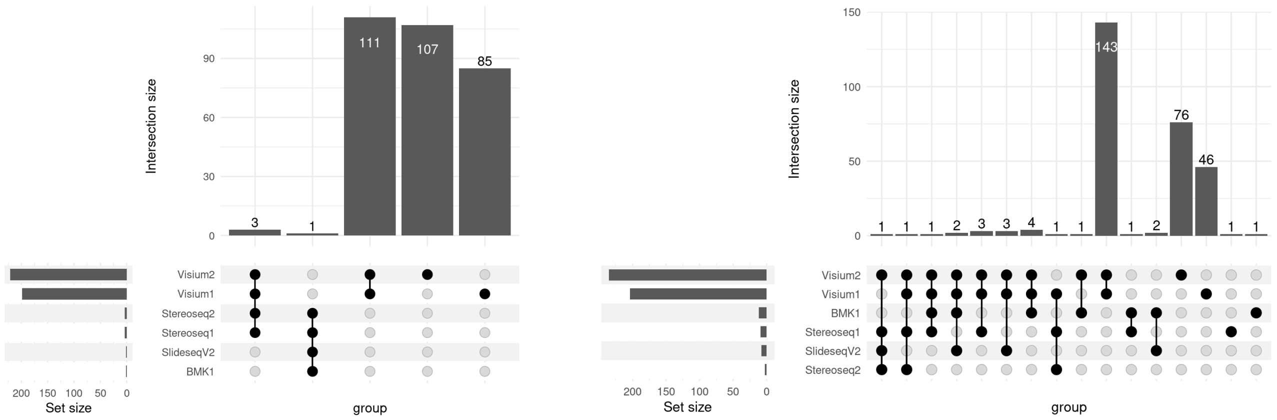


**Supplementary Figure 20**, Upset plots displaying the intersection of enriched ligand-receptor pairs obtained by different sST methods with results of *cellchat* shown in the left panel, results of *cellphoneDB* shown in the right panel.
